# VolumesJ: A new method and tool for volumetric estimation of brain structures after serial sectioning

**DOI:** 10.1101/2022.03.18.484850

**Authors:** Elisabete Ferreiro, Noelia Rodríguez-Iglesias, João Cardoso, Jorge Valero

## Abstract

Volume estimations are crucial for many neuroscience studies, allowing the evaluation of changes in the size of brain areas that may have relevant functional consequences. Classical histological methods and modern human brain imaging techniques rely on obtaining physical or digital sections, with a known thickness, of the organ to be analyzed. This “slicing” strategy is associated with an ineludible loss of information about the three-dimensional organization of the analyzed structures, especially affecting the precision of volumetric measurements. To overcome this problem, several methods have been developed. One of the most commonly used approaches for volume estimation is the classical Cavalieri’s method.

Within this book chapter, we provide first an overview of Cavalieri’s method and propose a new one, named the Truncated Cone Shape (TCS) method, for the estimation of volumes from tissue sections. Second, we compare the accuracy of both methods using computer-generated objects of different shapes and sizes. We conclude that, more frequently, the TCS method provides a better estimate of real volumes than Cavalieri’s method. And third, we describe a protocol to estimate volumes using a self-developed and freely available tool for ImageJ: VolumestJ (https://github.com/Jorvalgl/VolumestJ). This new tool helps to implement both Cavalieri’s and TCS methods using digital images of tissue sections. We consider that VolumestJ will facilitate the labor of researchers interested in volume estimations.

## 1 Introduction

Volume estimation of different brain regions is relevant in the context of neurodevelopment, aging, and pathology. Volume changes may indicate aberrant development under certain circumstances, such as in the presence of stressors, toxic agents, or inflammation [1–4]. Volume estimation can also help distinguish the moderate volume loss associated with some brain regions during normal aging from the large volume loss associated with age-related neurodegenerative diseases. [5, 6]. Additionally, estimates of volumetric cell densities obtained with unbiased stereological methods, such as the physical disector, may be combined with volume estimations to approximate the total number of cells in a given region of the brain [7].

The most commonly used microscopy visualization techniques of brain cells and structures require tissue sectioning, resulting in a subsequent loss of spatial (three-dimensional) information. Histological analysis of brain samples is a key method for neuroscience, and it has allowed a detailed description of the cellular organization of the central nervous system and the classification of their functionally relevant regions [8–11]. New methods, such as tissue clearing and light-sheet microscopy, have recently emerged, allowing imaging of whole brains and overcoming the necessity of sectioning [12]. However, these new methods come with their pitfalls, require a high time and economic investment, and are not widespread in the field. Therefore, tissue sectioning is still the most common technique for brain quantification. Additionally, the field of human brain clinical imaging utilizes thick brain section images. *In vivo* human brain imaging techniques, such as magnetic resonance imaging (MRI), produce low-resolution images (compared to microscopy) of thick virtual brain slices. Thus, both classical microscopy and human brain imaging confront the same problem when interpreting three-dimensional structures while looking at them in two dimensions. This problem was partially solved by stereology, defined by Davy and Miles [13] as “the science of inferring spatial structure from partial information, usually lower-dimensional data, which is usually in the form of sections or projections of the structure of interest”. However, despite the existence of several software tools to obtain stereological estimations, many neuroscientists still obviate the third dimension when performing cell counting or morphological measurements of brain regions and provide “surface measurements” or “area densities” when reporting morphological or cell number quantifications.

In this chapter, we will describe the classical Cavalieri’s method for volume estimation, and propose and validate a new method: the Truncated Cone Shape (TCS) method. Additionally, we describe a protocol to estimate volumes from digital images by using a self-developed and freely available ImageJ [14] tool: VolumestJ (for updated versions of the macro, please see https://github.com/Jorvalgl/VolumestJ).

## 2 Material

### 2.1 Generation of virtual objects

To validate and compare the accuracy of Cavalieri’s and TCS methods when estimating volumes from sectioned objects, we generated artificial (virtual) objects using the ImageJ package FIJI [15]. Virtual objects were created by generating consecutive sets of two-dimensional (2D) images which included white areas defining the sections of the virtual object. The different images of an object were concatenated forming one stack of images. Each section was considered to have 1 pixel in-depth, just for convenience as this is the minimal unit of a digital image. Therefore, each 2D image was considered to be separated from the consecutive next one by 1 pixel. The three dimensional (3D) superposition of these images (slices) formed the 3D virtual object. Different virtual objects were generated by concatenating a different number of images with different shapes. The properties of each virtual object are listed in Table 1.

**Table 1.** Properties of virtual objects used for validation (also check Figure 1).

| Name | Cavalieri's volume (pixels<sup>3</sup>) | TCS volume (pixels<sup>3</sup>) | No. Sections (1-pixel depth) | Max. width of 2D sections (pixels) | Max. height of 2D sections (pixels) |
| --- | --- | --- | --- | --- | --- |
| Sphere | 537314 | 537030.775 | 100 | 100 | 100 |
| Cylinder | 244800 | 243168 | 100 | 56 | 56 |
| Waved | 3291434 | 3279844.12 | 200 | 199 | 206 |
| Horn | 1000740 | 1000489.93 | 87 | 168 | 338 |

The Sphere was generated using a self-developed ImageJ macro (Macro code 1) that uses the formula of the section of the sphere (Equation 1) to create concatenated images of concentric circles of the adequate diameter *d* to create a sphere with a defined diameter *D*.

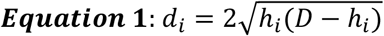

Where *d*_*i*_ is the **diameter of the circle** at the *i* slice that is located at a distance *h*_*i*_ of the surface of a sphere with diameter *D*.

The following macro code (Macro code 1) creates a sphere formed by 100 sections in a previously generated dark image. The number of sections may be altered by changing the value of the variable *planes*.

~~~
**Macro code 1**
//Sphere Generator Macro
/*
  Sphere_Generator is an ImageJ macro developed to generate spheres,
  Copyright (C) 2020 Jorge Valero Gómez-Lobo
  Sphere_Generator is free software: you can redistribute it and/or modify
  it under the terms of the GNU General Public License as published by
  the Free Software Foundation, either version 3 of the License, or
  (at your option) any later version.
  Sphere Generator is distributed in the hope that it will be useful,
  but WITHOUT ANY WARRANTY; without even the implied warranty of
  MERCHANTABILITY or FITNESS FOR A PARTICULAR PURPOSE. See the
  GNU General Public License for more details.
  You should have received a copy of the GNU General Public License
  along with this program. If not, see <http://www.gnu.org/licenses/>.
*/
//This macro has been developed by Dr Jorge Valero (jorge.valero@achucarro.org).
//If you have any doubt about how to use it, please contact me.
//License
Dialog.create(“GNU GPL License”);
Dialog.addMessage(“Sphere_Generator Copyright (C) 2020 Jorge Valero Gomez-Lobo.”);
Dialog.setInsets(10, 20, 0);
Dialog.addMessage(“Sphere_Generator comes with ABSOLUTELY NO WARRANTY; click on help button for details.”);
Dialog.setInsets(0, 20, 0);
Dialog.addMessage(“This is free software, and you are welcome to redistribute it under certain conditions; click on help button for details.”);
Dialog.addHelp(“http://www.gnu.org/licenses/gpl.html”);
Dialog.show();
planes=100;
for (i=1; i<=planes/2; i++){
          setSlice(i);
          d=2*(sqrt(i*(planes-i)));
          print(d);
          run(“Specify…”, “width=“+d+” height=“+d+” x=100 y=100 slice=1 oval centered”);
          setSlice(i);
          run(“Fill”, “slice”);
}
s=(planes/2);
for (i=planes/2; i>0; i--){
          s++;
          setSlice(s);
          d=2*(sqrt(i*(planes-i)));
          print(d);
          run(“Specify…”, “width=“+d+” height=“+d+” x=100 y=100 slice=1 oval centered”);
          setSlice(s);
          run(“Fill”, “slice”);
}
~~~

To run this macro code in FIJI, follow these instructions:

1. Create a new dark image with at least 100 slices and a recommended size of 200x200 pixels using the following sequence of tabs: File->New->Image.
2. Use the following sequence to open the Script console File->New->Script…
3. Copy the Macro code and paste it into the Script Console.
4. Select the ImageJ Language in the Language tab (Language->IJ1 Macro).
5. Run the macro by clicking the Run tab.
6. To get a three-dimensional view of the object use the 3D viewer plugin (Plugins->3D viewer).

The cylinder was generated manually by duplicating the same image of a white filled circle, and then concatenating all the images in a single stack with the ImageJ command concatenate (Image->Stacks->Tools->Concatenate…).

Random Waved object was generated by concatenating sections of different spheres with a random diameter. The images were generated by filling ROIs (regions of interest) with white pixels. The first ROI was a circle defined by us before running Macro code 2. Then, the macro randomly enlarged or decreased the area of the previous ROI and filled it to generate the next image. The stack of images was smoothed using the Gaussian Blur 3D filter of ImageJ, and the final result was converted into a binary image. The following macro code generates this type of random objects:

~~~
**Macro code 2**
//Random waved generator Macro
/*
   Random_waved_Generator is an ImageJ macro developed to generate a random waved 3D object,
   Copyright (C) 2020 Jorge Valero Gómez-Lobo
    Random_waved_Generator is free software: you can redistribute it and/or modify
   it under the terms of the GNU General Public License as published by
   the Free Software Foundation, either version 3 of the License, or
   (at your option) any later version.
   Random_waved_Generator is distributed in the hope that it will be useful,
   but WITHOUT ANY WARRANTY; without even the implied warranty of
   MERCHANTABILITY or FITNESS FOR A PARTICULAR PURPOSE. See the
   GNU General Public License for more details.
   You should have received a copy of the GNU General Public License
   along with this program. If not, see <http://www.gnu.org/licenses/>.
*/
//This macro has been developed by Dr Jorge Valero (jorge.valero@achucarro.org).
//If you have any doubt about how to use it, please contact me.
//License
Dialog.create(“GNU GPL License”);
Dialog.addMessage(“Random_waved_Generator Copyright (C) 2020 Jorge Valero Gomez-Lobo.”);
Dialog.setInsets(10, 20, 0);
Dialog.addMessage(“Random_waved_Generator comes with ABSOLUTELY NO WARRANTY; click on help button for details.”);
Dialog.setInsets(0, 20, 0);
Dialog.addMessage(“This is free software, and you are welcome to redistribute it under certain conditions; click on help button for details.”);
Dialog.addHelp(“http://www.gnu.org/licenses/gpl.html”);
Dialog.show();
setBatchMode(“hide”);
roiManager(“Select”, 0);
roiManager(“Add”);
for(i=100; i<200; i++){
        roiManager(“Select”, 0);
        run(“Enlarge…”, “enlarge=“+(−20*random)+(random*20));
        roiManager(“Update”);
        setSlice(i);
        fill();
}
for(i=100; i>0; i--){
        roiManager(“Select”, 1);
        run(“Enlarge…”, “enlarge=“+(random*-20)+(random*20));
        roiManager(“Update”);
        setSlice(i);
        fill();
}
run(“Select None”);
run(“Gaussian Blur 3D…”, “x=2 y=2 z=10”);
selectWindow(“Untitled”);
setAutoThreshold(“Default dark”);
setThreshold(1, 255);
setOption(“BlackBackground”, false);
run(“Convert to Mask”, “method=Default background=Dark”);
run(“Invert LUT”);
setBatchMode(“exit and display”);
~~~

To execute this macro a dark image with, at least, 200 slices and a recommended size of 500x500 pixels should be generated and a ROI of the desired shape added to the ROI Manager. To generate the ROI, please select the appropriate tool (eg. the Oval tool), draw the ROI in the image, and add it to the ROI Manager by clicking the key “t”. Then, run the macro following the instructions indicated for Macro code 1.

Finally, to generate the object named “Horn” we used a self-developed macro that takes an initial ROI, defined by the user, and generates a symmetric object. The macro generates ROIs with decreasing areas at both sides of the mid slice (slice 100) and fills them. To generate the “Horn” we drew a banana-like shaped initial ROI.

~~~
**Macro code 3**
//Symmetric object generator Macro
/*
    Symmetric_Object_Generator is an ImageJ macro developed to generate symmetric 3D objects,
    Copyright (C) 2020 Jorge Valero Gómez-Lobo
     Symmetric_Object_Generator is free software: you can redistribute it and/or modify
    it under the terms of the GNU General Public License as published by
    the Free Software Foundation, either version 3 of the License, or (at your option) any later version.
    Symmetric_Object_Generator is distributed in the hope that it will be useful,
    but WITHOUT ANY WARRANTY; without even the implied warranty of
    MERCHANTABILITY or FITNESS FOR A PARTICULAR PURPOSE. See the
    GNU General Public License for more details.
    You should have received a copy of the GNU General Public License
    along with this program. If not, see <http://www.gnu.org/licenses/>.
*/
//This macro has been developed by Dr Jorge Valero (jorge.valero@achucarro.org).
//If you have any doubt about how to use it, please contact me.
//License
Dialog.create(“GNU GPL License”);
Dialog.addMessage(“Symmetric_Object_Generator Copyright (C) 2020 Jorge Valero Gomez-Lobo.”);
Dialog.setInsets(10, 20, 0);
Dialog.addMessage(“Symmetric_Object_Generator comes with ABSOLUTELY NO WARRANTY; click on help button for details.”);
Dialog.setInsets(0, 20, 0);
Dialog.addMessage(“This is free software, and you are welcome to redistribute it under certain conditions; click on help button for details.”);
Dialog.addHelp(“http://www.gnu.org/licenses/gpl.html”);
Dialog.show();
roiManager(“Select”, 0);
roiManager(“Add”);
for(i=100; i<200; i++){
        roiManager(“Select”, 0);
        getStatistics(area, mean, min, max, std, histogram);
        run(“Enlarge…”, “enlarge=-1”);
        roiManager(“Update”);
        getStatistics(area2, mean, min, max, std, histogram);
        if (area==area2) i=200;
        else{
                 setSlice(i);
                 fill();
        }
}
for(i=100; i>0; i--){
        roiManager(“Select”, 1);
        getStatistics(area, mean, min, max, std, histogram);
        run(“Enlarge…”, “enlarge=-1”);
        roiManager(“Update”);
        getStatistics(area2, mean, min, max, std, histogram);
        if (area==area2) i=0;
        else{
                 setSlice(i);
                 fill();
        }
}
~~~

To execute this macro proceed as indicated for Macro code 2.

### 2.2. Automated analysis of virtual objects

To speed-up our analysis, we developed a strategy that automated the estimation of volumes of virtual objects. The previous macros generated black background and white foreground binary images, thus allowing easy automatic generation of ROIs by selecting all pixels with an intensity value above 0. With this automation capacity, we were able to analyze the volume of each object and to use different combinations of object slicing orientation, section thickness, and between section intervals. Therefore, we were able to check the behavior of Cavalieri’s and TCS methods in different conditions and with different error coefficients.

For each virtual object, the original image sections were further transformed to obtain many different stacks of section images combining three parameters: 1) three possible orthogonal slicing orientations, 2) different thicknesses, and 3) various section intervals. 1) For each virtual object, except for the sphere, we generated extra sets of image stacks by using the orthogonal sectioning tool of ImageJ (Image->Stacks->Reslice[/]…), avoiding interpolation and with an output spacing of 1 pixel. We used two different slicing planes: vertical and horizontal planes. 2) To simulate different section thicknesses, we fused consecutive images using the average projection from ImageJ: by fusing X images, we obtained sections with a thickness of X pixels. 3) On top of this, we also tested different between-sections intervals by using the stack reduce command of ImageJ (Image->Stacks->Tools->Reduce…). We designed a macro that iteratively generated and saved sequences of images with different thicknesses and section intervals (not provided, but available upon reasonable request) for subsequent analysis.

The final number of images generated *per* virtual object was large, and thus we created two versions of the macros included in the VolumestJ pack (Macro codes 4 and 5) that automated volume estimations of virtual objects. First, we developed a fully automatic version of Macro code 4, which is available upon reasonable request. This macro generated ROIs by selecting all pixels above threshold level 0 and saved them with the adequate organization and format to be subsequently processed with an automatic version of VolumestJ. Second, we created a special version of VolumestJ (Macro code 5) that automatically analyzed all the stack of images generated from each virtual object and saved area, estimated volumes, and error data (not provided, but available upon reasonable request).

### 2.3 VolumestJ testing with real images from the mouse hippocampus

We tested the utility of our TCS method and VolumestJ to identify previously described sexual dimorphism in hippocampal volume, by estimating the volume of a specific region of the mouse hippocampus in female and male mice: the septal portion of the granule cell layer of the dentate gyrus (GCL). We defined the septal hippocampus as the most dorsal part of the hippocampus spanning from bregma -1.70 to -2.3 mm in the anteroposterior axis. Full details of animals and tissue processing and fluorescent microscopy imaging are provided in the following sections.

### 2.4. Animals

All experiments were performed in 3.5 month-old female (n=6) and male (n=7) B6.Cg-Tg(Csf1r-EGFP) mice [16]. Mice were housed in a 12:12 hrs light cycle, with ad libitum access to food and water. Mice were anesthetized with an intraperitoneal administration of Avertine (2,2,2,-tribromoethanol, 0.250 mg/g), and perfused with Phosphate Buffer Saline (PBS) to rinse the blood, followed by 4% paraformaldehyde in PBS (PFA, 1.5 ml/g). Finally, brains were dissected out, post-fixed for 3 h in 4% PFA, and stored in an anti-freezing solution (30% v/v sucrose, 30% v/v ethylene glycol in deionized water) at -20°C until sectioning. All procedures followed the European Directive 2010/63/EU and were approved by the Ethics Committees of the University of the Basque Country EHU/UPV (Leioa, Spain; protocol reference: CEEA_M20_2019_279).

### 2.4 Brain sectioning

Mouse brains were divided into the right and left hemispheres, and the cerebellum and olfactory bulb of the left hemispheres were removed. Brain blocks were glued to the mobile platform of the LeicaVT1200S vibrating blade microtome (Leica Microsystem GmbH, Wetzlar, Germany) with a thin layer of cyanoacrylate. 50 μm-thick sagittal sections were obtained using a vibrating amplitude of 1.2 mm and a blade forward speed of 1.2 mm/s. Brain slices were collected in PBS and divided into 6 series (6 wells of a 24 multiwell plate). One of the six series was used for volume estimations, meaning that we used a between-sections interval (Si) of 1/6. Non-used tissue sections were stored in the anti-freezing solution at -20°C.

### 2.5 Immunofluorescence staining

Tissue sections were rinsed in PBS (3x10 min) at room temperature, with constant shaking, then blocked in permeabilization solution (0.2% Triton X-100, 3% bovine seroalbumin in PBS; all from Sigma-Aldrich) for 1 h. Tissue sections were used to detect different epitopes for other research projects. Thus, they were incubated in the permeabilization solution with primary antibodies for 48 hrs at 4°C with constant shaking. Then, slices were rinsed in PBS (3x10 min) at room temperature and with constant shaking. Tissue sections were incubated with secondary antibodies, and 2-(4-Amidinophenyl)-6-indolecarbamidine dihydrochloride (DAPI, 1:1000, D9542, Sigma Aldrich), which we regularly use to stain nuclei, diluted in permeabilization solution for 2 hrs at room temperature and with constant shaking. Finally, sections were rinsed with PBS (3x10 min) and mounted on glass slides with Dako Cytomation Fluorescent Mounting Medium (S302380-2, Agilent Technologies).

### 2.6 Microscopy imaging

Fluorescence images from each slide were collected using a 3DHistech Pannoramic MIDI II Slide Scanner (3DHISTECH Ltd.). We used a Plan-Apochromat 20x objective (NA: 0.8) and a camera adapter with a 1.6x magnification index. The resolution of the raw images was 0.203125 μm^2^/pixel, but images were converted to tiff format and downsized for analysis to a final resolution of 1.6250 μm^2^/pixel.

### 2.7 Statistical analysis

Data from virtual objects were analyzed using GraphPad Prism 5 (GraphPad Software, Inc, San Diego, CA) to check differences i regression lines (all of them with a slope significantly different from 0: *p*<0.0001) obtained for two different types of calculated errors (equations 4 and 12) for Cavalieri’s or the TCS methods. Only *p*<0.05 was considered significant. We also compared error differences between Cavalieri’s method and the TCS using DABEST software [17]. Error pairs, obtained for the same stack of images of a determined virtual object (with the same slicing orientation, thickness, and section interval), were analyzed using the median paired differences. Data from mouse hippocampi were also analyzed using DABEST software to calculate mean differences. The *p*-values reported are the likelihoods of observing the effect sizes if the null hypothesis of zero difference (median paired or mean) is true. For each permutation *p*-value, 5000 reshuffles of the control and test labels were performed. Median and mean differences were plotted as bootstrap sampling distributions and depicted as dots. The ends of the vertical error bars in the plots indicated 95% confidence intervals. To compare the variability obtained with the different methods when analyzing GCL volume estimations, we calculated the relative standard error (rse). The rse is the percentage that represents the standard error of the mean (sem) with respect to the total mean (sem/mean*100).

## 3 Methods

### 3.1 Cavalieri’s method for volume estimations

Cavalieri’s estimator has been widely used in stereology to infer volumes from bi-dimensional images [18–21]. The main advantage of this estimator is that it does not require any assumption about the shape of the three-dimensional object, or brain region, to be analyzed. The “only” requirement is the use of systematic and adequate sampling, which is not trivial [18, 22–24].

Two are the key sampling steps for Cavalieri’s estimation method: 1) the selection of sections in which area estimations will be performed, and 2) area sampling of the region of interest in each particular section. Equidistant and similarly oriented sections are selected in the most common version of Cavalieri’s method. The thickness of the sections is known, while the area of the region of interest inside the sections is what needs to be estimated. Additionally, the interval between selected sections directly affects the accuracy of the method, and it may be chosen empirically considering the variability in shape and size of the region of interest along the cutting axis. Coefficients of error (CE) associated with the variability in the shape and size of the regions studied may be used to empirically evaluate the accuracy of the method [22, 25]. We will expand the concept of CE at the end of this section.

Classically, Cavalieri’s estimation of volumes is preceded by the estimations of the regions of interest inside the selected sections. To facilitate unbiased estimation of areas, area sampling is implemented by superimposing a point counting grid to the images of tissue sections. Each point in the grid is equidistant from the surrounding ones and represents a determined real tissue area or, if the thickness of the tissue slice is considered, a determined real volume. The size of the area/volume represented by each counting point will depend on the density of points in the grid. Thus, the higher the density of the grid, the better the accuracy of the estimated area. Nowadays, the advantages offered by digital imaging allow the use of the minimum unit of a digital image as the counting point of the grid: pixel or voxel. Each pixel/voxel of the image has an associated area/volume, which is already embedded into the metadata of calibrated images. Thus, by drawing the contour of a determined area in the digital image, it is possible to automatically and immediately infer its associated area/volume based on pixel/voxel size. Using ImageJ’s polygon tool, only if at least half of a given pixel is within the drawn contour will it be included in the estimates. Considering this, counting grids for volume estimation has become obsolete as it does not offer a real advantage over pixel-based size estimations. Nevertheless, it should be considered that image resolution becomes a new parameter to be considered when designing our quantification protocol, as it will directly affect the accuracy of the estimation method.

Cavalieri’s method uses the following equation to estimate **volumes** [18]:

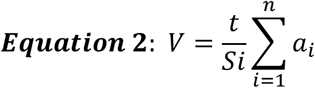

Where *S*_*i*_ is the section interval (eg. 1/6 when 1 in 6 sections are selected for quantification), t is the mean section thickness, *n* the number of sections, and *a*_*i*_ is the estimated area of section *i*. The **area of each section** is estimated using equation 3:

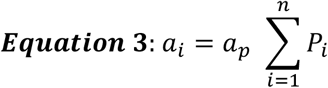

Where *a*_*p*_ is the area associated with each point of the grid or with each pixel in digital images, and *P*_*i*_ is the number of points or pixels enclosed by the area of interest in a determined section *i*.

Importantly, Cavalieri’s estimations have associated **coefficients of error** (m0 CE and m1 CE), which may be calculated using the following equations [22, 25]:

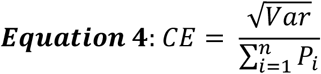

Where *Var* is the **total estimated variance**, which is the sum of the noise variance (*S*^2^), and the variance associated with systematic random sampling (*Var*_*SRS*_):

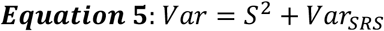

The **variance due to the noise** is calculated using the following equation:

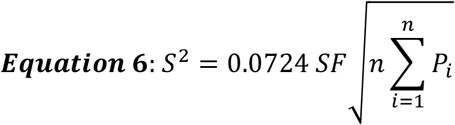

Where *SF* is the **shape factor** and it is calculated as the ratio between the perimeters and the squared root of the area of the analyzed regions.

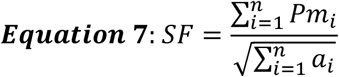

Where *Pm*_*i*_ is the perimeter of the region of interest in each section *i*.

Finally, *Var*_*SSS*_ for m0 CE (*Var*_*SRSm0*_) and m1 CE (*Var*_*SRSm*1_) are calculated using the following equations:

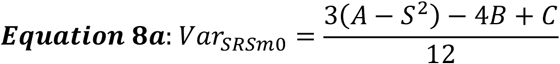

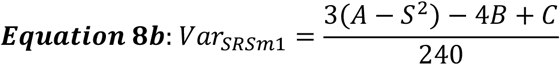

Here, it is possible to use two different denominators, 12 or 240, which depends on considerations about the size of the space between consecutive sections [23]. When using *Var*_*SRSm*0_, we will here name the CE as m0 CE, and when using *Var*_*SRSm*1_, the name m1 CE will be used. We have always used m1 CE in our studies, as the distance between consecutive sections used in microcopy is very small. A, B, and C are calculated as follows:

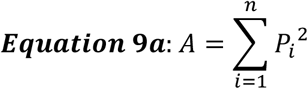

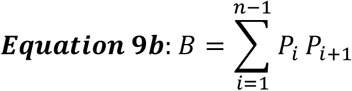

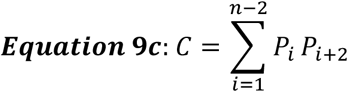

When performing volume estimations, it is highly recommended to perform pilot studies using different section intervals (*Si*) and image resolutions (the modern equivalent to the old counting grid size that will determine the number of points counted *per* section: *P*_*i*_). This will provide an estimate of the CE associated with each particular strategy of quantification. Then, the results obtained in the pilot study will serve to select the best protocol parameters, mainly image resolution and section interval, for our quantification protocol. This protocol design is essential to save time and resources while obtaining good enough accuracy of the measurements. We here reviewed the validity of the CE, by comparing it with the real accuracy of volume estimations of virtual objects of known volumes (section 3.3).

### 3.2 Truncated Cone Shape (TCS) method for volume estimations

In Cavalieri’s method, the areas of omitted sections are considered equal to the areas of the analyzed ones. Therefore, this approximation does not consider the natural smooth variations in the area expected for sequential sections of biological samples (graphs in Figure 1). Thus, depending on the shape of the structure and the fraction of sections analyzed, Cavalieri’s estimations may present some inconveniences compared with other estimators that consider smooth variations between the areas of adjacent sections.

**Figure 1.**
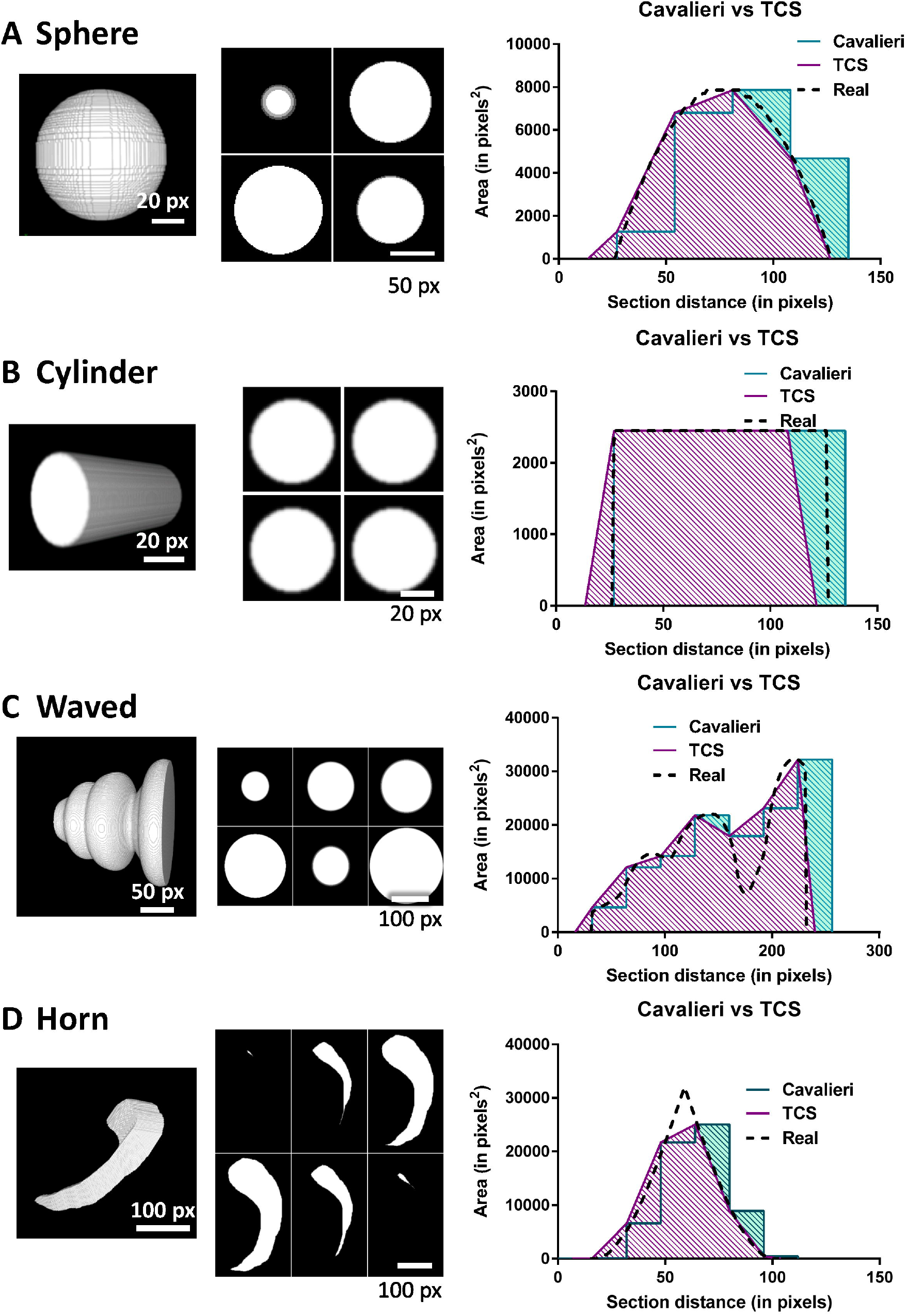
Virtual objects for TCS method validation. **A, B, C, and D** show three-dimensional renderings (left images) of the virtual objects created (left column images), an example of the two-dimensional images generated to create virtual objects (mid-column images), and the curves obtained by representing changes in section areas along the slicing axis. The area under these curves represents the volume. “Real” refers to the curve obtained using all the images generated to create the virtual objects.

Previously, we used sophisticated software (such as TableCurve 2D, by Systat Software Inc) to fit the changes in the area of the sections along a determined slicing axis to a curve function. The integral of this function allowed the estimation of the volume: the area under the curve [26, 27]. However, this method has the inconvenience of requiring a high level of expertise for the selection of the adequate curve function which better fits the analyzed data. As a relatively simple and reliable new stereological option, we propose here a “truncated cone shape” approach. This method estimates volumes considering: 1) that the area of the analyzed region in each section may be re-interpreted as the area of a circle, and thus 2) that the volume between two consecutive measured sections is approximately equivalent to the volume of a truncated cone. In this cone, the base and top areas are equal to the areas of two consecutive measured sections, and the distance between the sections equals the height of the truncated cone. Therefore, the **total volume** may be calculated as the sum of the volumes of the truncated cones defined between each consecutive pair of measured sections using the following equation:

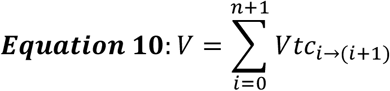

Where *n* is the number of sections and *Vtc*_*i*−*i*+1_ is the estimated volume of truncated cones defined between each pair of consecutive sections *i* and *i+1*. For our estimations we have included two not measured sections, sections 0 and *n+1*, situated at a distance equal to half of the regular distances between consecutive sections and with a volume of 0. This strategy smooths the ends of the volume avoiding abrupt borders.

Each **volume** is calculated using the following equation:

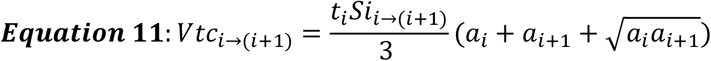

Where *t*_*i*_ is the thickness of section *i, Si*_*i*→*i*+1)_ the sections interval between sections *i* and *i+1*, and *aa*_*i*_ the estimated area of section *i* (Equation 3). Thus, this equation admits different section intervals and thickness. However, considering that adequate sampling may be relevant, we recommend the use of regular thickness and section intervals.

The same error equations used for the Cavalieri’s method (equations 3 to 8) may be used with the TCS method as the total variance is also due to both noise variance (*S*^2^) and the variance associated with systematic random sampling (*Var*_*SRS*_). However, it should be noticed that the precision of the TCS method is highly dependent on the proper ordering of the sections, something that is not relevant when using Cavalieri’s estimation.

### 3.3 Comparison of Cavalieri’s and TCS methods for volume estimations

Cavalieri’s method has been deeply revised, and it is considered the gold standard method for volume estimations. However, the TCS method is less extended in scientific and clinical imaging. Methods similar to the TCS have been employed to determine the volume of various body parts, as well as the whole body in humans [28, 29] and other mammals [30]. Additionally, this approach has been used to estimate the volume of human muscles [31] and pathological structures such as atherosclerotic plaques [32], lymphedema effects on limbs [33], and tumors [34, 35].To our knowledge, we have been the only ones applying it for volume estimations in neuroscience papers [36, 37]. To be able to compare and validate the new method, we have generated four artificial objects with different morphologies and known volumes (Table 2 and Figure 1). For the evaluation, we simulated artificial slicing of the objects with different thicknesses in three orthogonal planes as described in section 2.2. We also used different section intervals, being 2 the minimum number of sections per object. The combination of a determined section thickness with a specific section interval constituted one trial. A representative trial *per* object is shown in Figure 1. Trials in Figure 1 have been selected by their similar CEs (m1 CE close to 0.02). For each trial we were able to calculate the **percentage of error of the method** using the following formula:

**Table 2.**
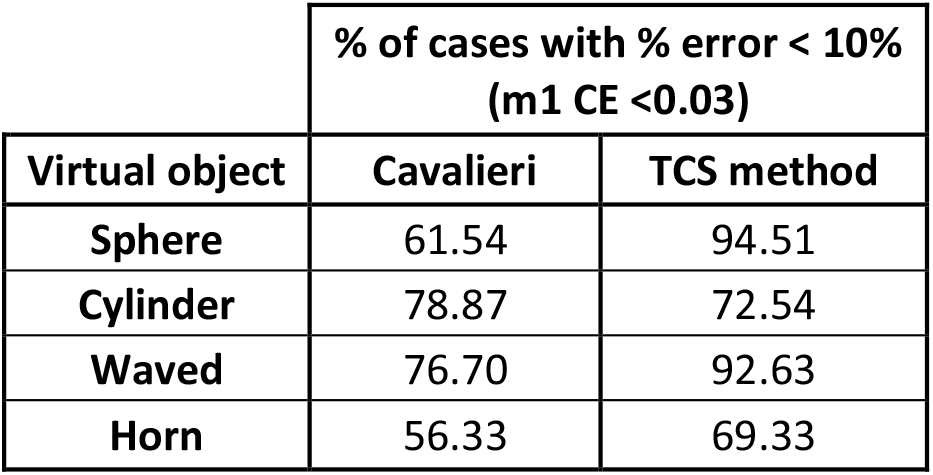
The proportion of volume estimations with an m1 CE below 0.03 and a % of error below 10%.

| % of cases with % error < 10% (m1 CE <0.03) |  |  |
| --- | --- | --- |
| Virtual object | Cavalieri | TCS method |
| Sphere | 61.54 | 94.51 |
| Cylinder | 78.87 | 72.54 |
| Waved | 76.70 | 92.63 |
| Horn | 56.33 | 69.33 |

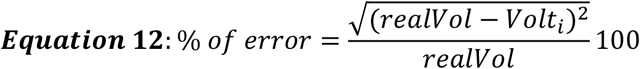

Where *realvol* is the volume obtained using the maximum number of sections available (thickness=1 pixel, and *Si*=1/1 section) with the method to be evaluated, and *Volt*_*i*_ is the volume obtained in trial *i* with the same method.

We analyzed all the trials with a minimum number of two sections using linear regression models depicted in Figure 2A, C, E, and G. From the four simulated objects analyzed, only the analysis of the cylinder did not show significant differences between both methods when checking the general trend of the % of error with the linear regression. The sphere and horn showed a lower error regression line for data obtained with the TCS method. The waved shaped object showed an equal or lower error regression line using the TCS method mainly for combinations of pairs of section thicknesses and intervals that led to an m1 CE bellow 0.03 (Figure 2). Curiously, Cavalieri’s method showed a better performance when estimating the volume of the waved object for m1 CE values above 0.03. Additionally, we detected a general better correspondence between the m1 CE and the % of error with the TCS method, except for the horn object, as indicated by the determination coefficient of the regression lines: r^2^. This suggests that the m1 CE is as valid as an indicator of the accuracy of the TCS method as it is for Cavalieri’s method. To complement these results, we also analyzed the proportion of section thickness and interval pairs that gave a m1 CE bellow 0.03 and showed a % of error below 10% (Table 2). Our data indicate that the TCS method is, except for the cylinder, more accurate than Cavalieri’s method. We also specifically analyzed significant differences between the % of error of the two methods used. Again, only for the cylinder, Cavalieri’s method showed a lower error, while the TCS method was more accurate for the other three objects (Figure 2B, D, F and H).

**Figure 2.**
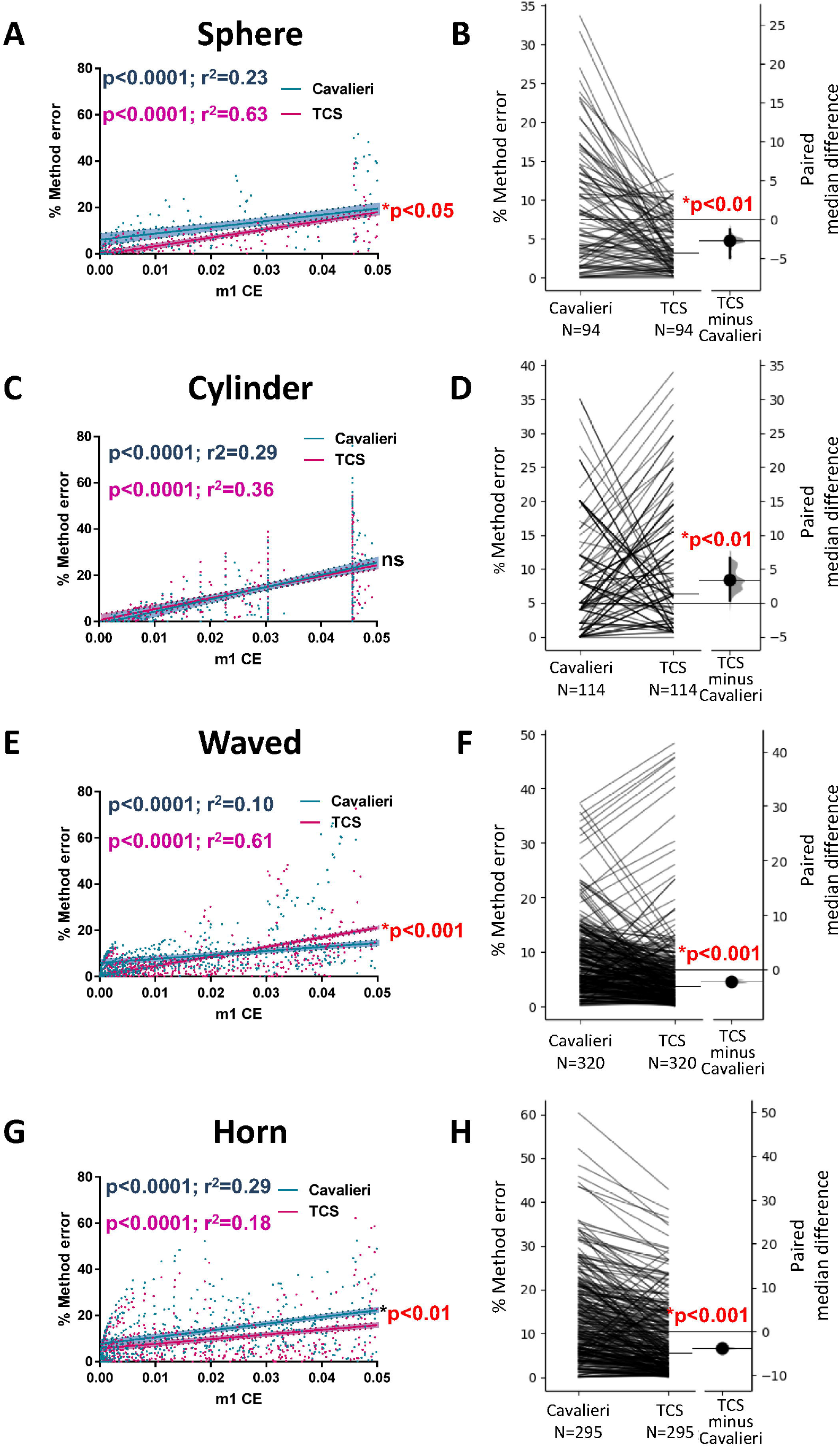
Validation of Cavalieri’s and TCS methods. **A, C, E, and G** show the regression lines obtained when confronting the % of error of each method (Equation 12) with their corresponding m1 CE (Equation 4). Dots correspond to individual estimations from a specific set of images representing a particular pair of section thicknesses and intervals. Dotted lines and filled areas around regression lines represent the 95% confidence interval. Cyan and magenta *p-* values indicate slopes significantly different from 0, while red *p-*values indicate significantly different slopes between Cavalieri’s and TCS method regression lines. The coefficients of determination (r^2^) indicate that for most objects (except for the Horn), the TCS method shows a better correspondence between the calculated error and the m1 CE. This confirms that m1 CE is also a valid estimator of the accuracy of the TCS method. **B, D, F, and H**. Gardner-Altman estimation plots show the paired median difference between the % of error of Cavalieri’s and the TCS methods. Both groups are plotted on the left axes as a slope graph: each paired set of observations is connected by a line. The paired mean differences are plotted on floating axes on the right as bootstrap sampling distributions. The mean differences are depicted as dots; the ends of the vertical error bars indicate the 95% confidence intervals. The *p*-values are the likelihoods of observing the effect sizes if the null hypothesis of zero difference is true. For each permutation *p*-value, 5000 reshuffles of the Cavalieri’s and TCS method labels were performed. Cavalieri’s method error is significantly smaller only for the cylinder, while for the other three objects the accuracy of the TCS method is significantly better.

Summarizing, the TCS method provides a better estimation of the real volume of the analyzed objects except for the cylinder. The cylinder presents a main differential characteristic when compared to the other objects: the shape and area of the sections do not vary along the slicing axis. Therefore, is it very likely that Cavalieri’s method shows a similar or even better performance than the TCS method when estimating the volume of 3D structures in which the sections are similar in size and shape, such as any type of geometrical prism. Based on the results obtained here, we suggest the use of the TCS method in combination with an adequate sampling strategy: the combination of section thickness and interval that gives a good enough m1 CE (at least below 0.03).

### 3.4 VolumestJ: A new freely available tool to estimate volumes using Cavalieri’s and TCS methods

Considering the relevance of analyzing volumes and with the expectation of facilitating the future labor of researchers aimed to perform stereological estimations, we have developed a pack of two ImageJ macros which will allow stereological estimation of volumes using both Cavalieri’s and TCS methods (freely available at https://github.com/Jorvalgl/VolumestJ). The first macro (VolumestJ_ROIsGenerator) is used to define the ROIs to be measured and is needed to appropriately save them and organize images for the next step in the analysis. The second macro (VolumestJ) automatically opens ROIs and Images, saving the areas, pixels, and perimeters of the ROIs in tables. After this, the macro asks the users to check the correct order of the sections, allowing them to change their order. Finally, the macro estimates volumes using Cavalieri’s and TCS methods based on equations 2 and 3, and 10 and 11 respectively. Besides, the CEs (both m0 and m1) are also calculated using equations from 4 to 9.

#### 3.4.1. Macro installation and running

For image analysis, we recommend the usage of the ImageJ package FIJI (ImageJ 1.53F) [15] in which our tool was developed. Our ImageJ macros may be installed using two different methods:

1. Installation of the macros as an ImageJ plugin:
  1.1) download the files from the supplementary material of this chapter,
  1.2) unzip the files,
  1.3) copy macro files (ended with .ijm) and paste them in your computer’s ImageJ plugins folder,
  1.4) start or restart FIJI; the macros should appear in the Plugins menu. Click on their name to run them.
2. Running macros from ImageJ script window:
  2.1) generate a new script in FIJI (File->New->Script…),
  2.2) activate the ImageJ Macro Language in the Script window (Language->IJ1 Macro),
  2.3) copy and paste the Macro code (Macro code 4 and Macro code 5, see below) in the Script window,
  2.4) and run the Macro (Run tab).

The first step of the analysis is the generation of the ROIs. The macro VolumestJ_ROIsGenerator should be used for this purpose. This macro will assist the user during the manual delineation of ROIs in each section. It will open the images, ask the user to draw ROIs, and save the ROIs in the appropriate folder with a name code related to the name of the image, thus allowing the correct functioning of the next macro (VolumestJ).

#### 3.4.2. Step by step guide of use for VolumestJ_ROIsGenerator

1) Immediately after running the macro, a window with the Copyright License disclaimer will appear. To continue just click OK (more information about this License may be found by clicking the Help button).
2) A new window will prompt asking the user to select the folder that contains the images. Select the adequate folder and click OK. The folder should contain subfolders allocating the images of individual specimens (ie. one subfolder *per* experimental or control case, Figure 3). We have developed the macro to be able to work with images containing one or more sections (eg. images from a slide scanner). At this point, the macro will automatically create two folders: the Processed folder, where processed images (ie. those images for which ROIs have been already generated) will be moved, and the ROIs folder, where newly-generated ROIs will be saved.
3) A window asking for the number of different ROI types to be generated will prompt (Figure 4A). Here, the user should indicate the number of different structures (eg. brain regions) to be analyzed *per* specimen, sample, or object. Therefore, it is possible to estimate the volume of different regions contained in the same group of images by indicating a number above 1 in this window. This involves that more than one type of ROI should be generated *per* image in the next step. To better illustrate this, we will use an example. To quantify the volume of the whole hippocampus (ROI 1) and a specific layer inside the hippocampus (ROI 2), the granular cell layer (GCL), we should write 2 in this window and click OK.
4) A parameters menu will prompt asking for three different types of information (Figure 4B): a. the name of the ROIs (predefined names will appear as ROIx, where x is the number of the ROI), b. the channel of the image where the user will draw the ROIs, b. and a checkbox to indicate whether and automatic contrast enhancement should be applied to the images. When checking the Enhance contrast box the macro will apply the Enhance contrast command of FIJI (Process->Enhance Contrast…) saturating 35% of the pixels of the image. This will normally allow better visualization of the images but they will not be saved with this new contrast, and original images will be preserved. After clicking OK the selected parameters will be saved in a .txt file named ROIGParameters.txt in the same directory that contains the Image folder. The .txt file will be used in other sessions to avoid steps 3 and 4. If the user wants to use new parameters, just remove or reallocate the ROIGParameters.txt file.
5) The first image will be opened, and a new window will prompt asking the user to draw ROIs (Figure 4C). The user should delineate the region to be analyzed and add them to the ROI Manager by clicking t or clicking in the Add tab of the ROI Manager (ROI Manager window may be opened using the following sequence of ImageJ tabs: Analyze->Tools->ROI Manager…). If the region to be delineated appears fragmented in the image, just draw and add the necessary number of ROIs to the ROI Manager, the next macro will detect more than one ROI for this specific region and fuse them before obtaining area estimations. Once the ROI or ROIs have been added to the ROI Manager, click OK and the macro will continue. If you added a ROI to the ROI Manager it will continue with step 6a. If the image does not contain the region to be analyzed just click OK without adding any ROI to the ROI Manager and the macro will proceed with step 6b.
6a) If ROIs have been added to the ROI Manager, a new window will prompt asking the user to confirm whether she/he wants to save the ROI or ROIs shown (Figure 4D). By clicking the tab Yes the macro will save the ROI or ROIs to the ROIs folder and continue to the next step. By clicking tab No, the macro will repeat step 5; thus allowing the user to change, delete or add new ROIs. The Cancel tab will end the macro. It is important to notice that if the macro is cancelled at this point the last ROI will not be saved.

**Figure 3.**
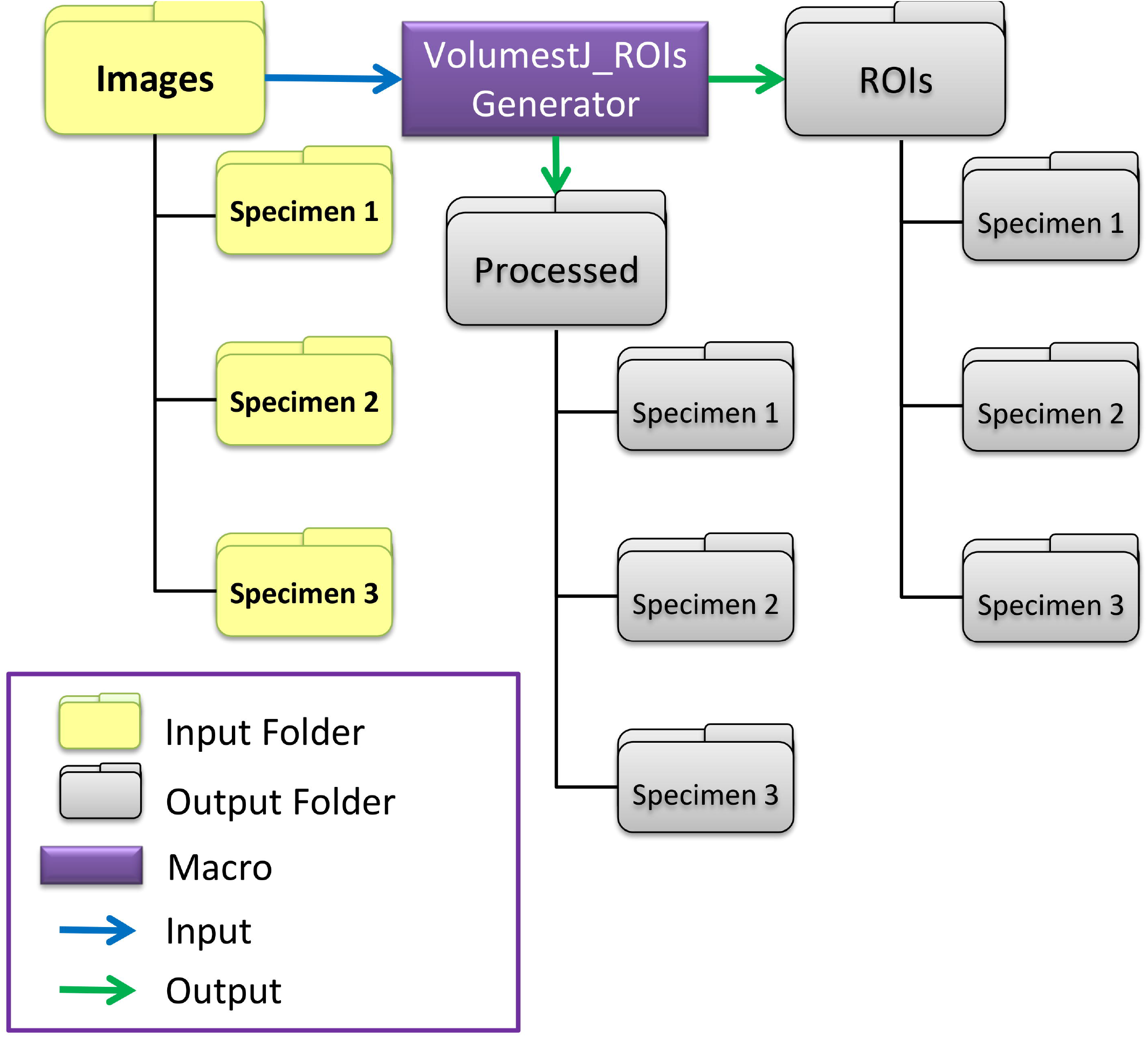
Folder organization requirements for ROIs generation. VolumestJ_ROIsGenerator macro requires images from the same object or specimen to be grouped into subfolders located inside a more general folder. Notice that, in the scheme, we named this general input folder Images for clarity reasons, but the name is irrelevant. The user should select this general Images folder when running the macro. The macro will generate output folders moving the images to a folder named Processed. Moved images will be organized in the same subfolders found in the Images folder. ROIs will be saved to the ROIs folder and organized following a subfolders structure similar to the Images folder.

**Figure 4.**
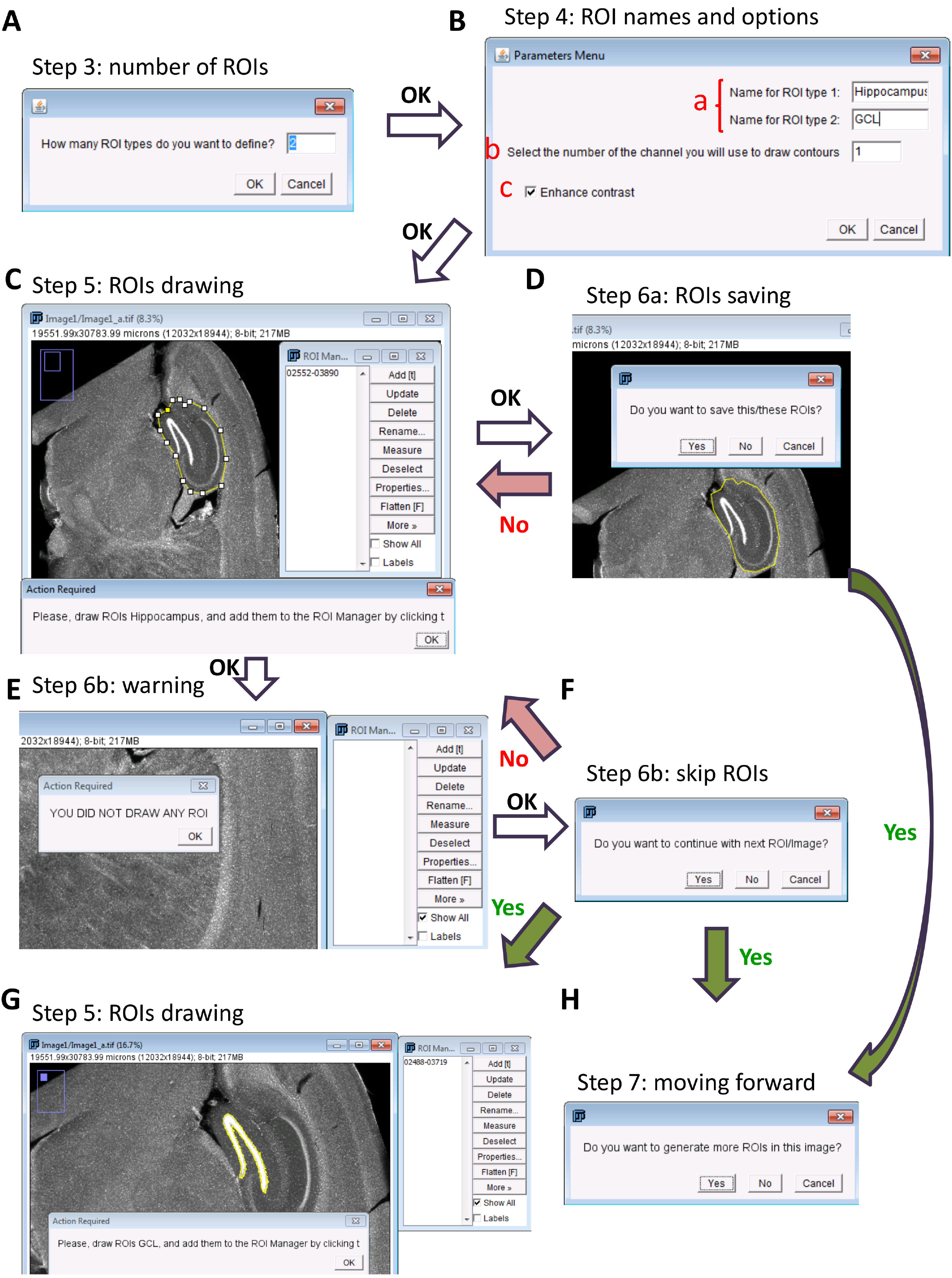
Relevant steps for ROIs generation with VolumestJ_ROIsGenerator. VolumestJ_ROIsGenerator macro helps users generate ROIs to estimate volumes with VolumestJ. **A**. Step 3 pop up window asking for the number of different types of ROIs to generate. **B**. After clicking OK in step 3 this menu window will prompt (step 4), where the user should indicate: a. the name of the ROIs, b. the channel she/he will use to draw ROIs, and c. if image contrast should be enhanced (this will generally allow a better visualization). **C**. Clicking OK in the window depicted in **B** the macro will proceed to step 5 opening an image, and the user will be asked to draw the ROIs of one of the types specified in **B** (step 4). ROIs should be added to the ROI Manager by clicking t or the Add button. More than one ROI may be drawn at this step if the area to be measured appears in more than one region of a section. Once all the ROIs of the required type associated with a single section have been added to the ROI manager, click OK, and the macro will proceed to step 6a (**D**). Additionally, if the region of interest does not appear in the image, just click OK without drawing any ROI. Then, the macro will detect the absence of ROIs and will proceed to step 6b (E). D. If at least one ROI was added to the ROI manager the macro will detect it allowing the user to save it and proceed to the next step (G or H) by clicking Yes, or to go back to step 5 (C) by clicking No (this allows the user to correct mistakes made in step 5). E. When the macro does not detect any ROI in the ROI manager it will proceed to step 6b showing a warning message. **F**. After clicking OK, a new window will prompt asking the user to decide if proceed to the next step (**G or H**) and skip the drawing of ROIs in the current section by clicking Yes, or to go back to step 5 (**C**) having the opportunity of drawing a new ROI. **G**. The macro will repeat step 5 to 6 until all types of ROIs defined in step 4 (**B**) have been saved or skipped. **H**. Once all the ROI types have been saved or skipped the macro will ask if the user wants to draw more ROIs in the current image. This is useful if more than one section appears in the same image. By clicking Yes the macro will proceed to step 5 (**C**), maintaining the current image opened. If No is clicked, the macro will close the current image, open the next one, and proceed to step 5 (**C**).
6b) If the user did not add any ROI to the ROI Manager a warning message will appear indicating that no ROIs were added (Figure 4E). After clicking OK a new window will appear asking the user to make a decision (Figure 4F), whether to proceed with the next step without saving any ROI (by clicking Yes tab) or to repeat step 5 (by clicking No tab). The Cancel tab will end the macro.
7) The macro will repeat steps 5 to 6 until all the ROI types have been generated (Figure 4G) and then ask the user whether she/he wants to generate more ROIs in the current image (Figure 4H). This is useful when using images that contain more than one section. In this case, ROIs should be created individually for each section. By clicking Yes, the macro will repeat step 5 without opening a new image. By clicking No the macro will close the current image, move it to the Processed folder, and repeat step 5 opening a new image. The macro will end once the Images Folder is empty. If the user wants to generate ROIs in an already processed image, just move back the image from the Processed folder to the original Images Folder and run the macro.

The final output of VolumestJ_ROIsGenerator will be two new folders: Processed and ROIs (Figure 3). The Processed folder will contain original images allocated in their corresponding subfolders, and the ROIs folder the newly-generated ROIs allocated in their corresponding subfolders. Additionally, subfolders for each ROI type will be also generated (Figure 5). The name of each ROI will contain the name of the image without the extension, then a number corresponding to the order of generation, and finally the extension .zip. This coding of ROI names is an essential requirement for their use with the VolumestJ macro.

**Figure 5.**
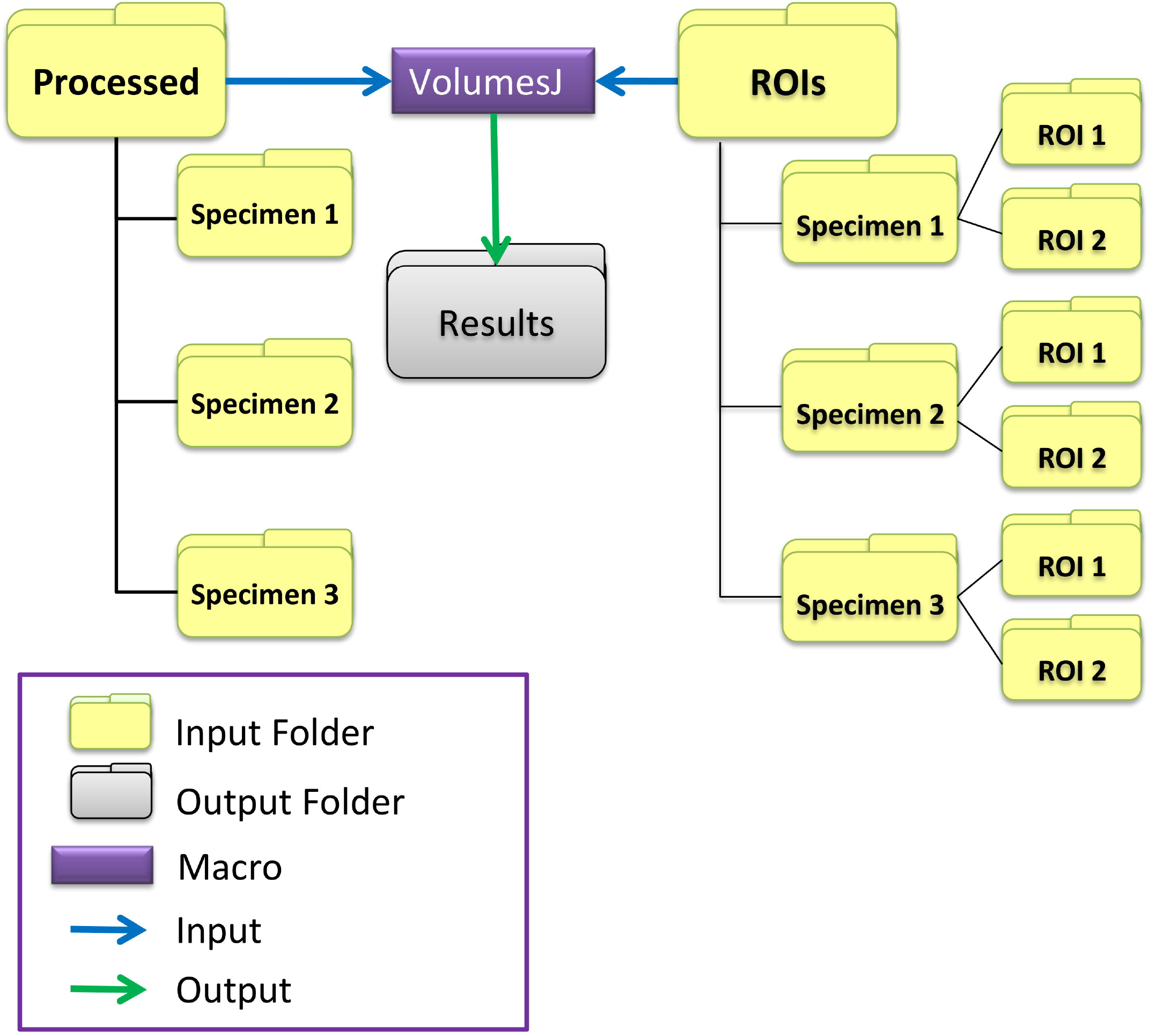
Folder organization requirements for volume estimations with VolumestJ. VolumestJ macro requires the existence of the Processed and ROIs folders (capital letters in the names are relevant). The Processed folder must contain images from the same object or specimen grouped into subfolders. The ROIs folder must contain ROIs from the same object or specimen grouped into subfolders, and these subfolders should be named equal to those contained in the Processed folder. ROIs subfolders must contain one folder *per* ROI type with the ROIs files. ROIs files must be named with the name of their corresponding image without the extension, then a number corresponding to the generation order, and ending in .zip. Therefore, we recommend using VolumestJ_ROIGenerator macro to create ROIs, as this macro will organize images and ROIs matching the requirements described here. VolumestJ macro will create a new output folder named Results containing .xls files with all data tables generated (ie. pixel, area, perimeter, and volume tables).

~~~
**Macro code 4**
*// VolumestJ_ROIsGenerator_v1.2.ijm
/*
   VolumestJ_ROIsGenerator is an ImageJ macro developed to manually generate ROIs,
   Copyright (C) 2020 Jorge Valero Gómez-Lobo
   VolumestJ_ROIsGenerator is free software: you can redistribute it and/or modify
  it under the terms of the GNU General Public License as published by
   the Free Software Foundation, either version 3 of the License, or
   (at your option) any later version.
   VolumestJ_ROIsGenerator is distributed in the hope that it will be useful,
   but WITHOUT ANY WARRANTY; without even the implied warranty of
   MERCHANTABILITY or FITNESS FOR A PARTICULAR PURPOSE. See the
   GNU General Public License for more details.
   You should have received a copy of the GNU General Public License
   along with this program. If not, see <http://www.gnu.org/licenses/>.
*/
//This macro has been developed by Dr Jorge Valero (jorge.valero@achucarro.org).
//If you have any doubt about how to use it, please contact me.
//License
Dialog.create(“GNU GPL License”);
Dialog.addMessage(“VolumestJ_ROIsGenerator Copyright (C) 2020 Jorge Valero Gomez-Lobo.”);
Dialog.setInsets(10, 20, 0);
Dialog.addMessage(“VolumestJ_ROIsGenerator comes with ABSOLUTELY NO WARRANTY; click on help button for details.”);
Dialog.setInsets(0, 20, 0);
Dialog.addMessage(“This is free software, and you are welcome to redistribute it under certain conditions; click on help button for details.”);
Dialog.addHelp(“http://www.gnu.org/licenses/gpl.html”);
Dialog.show();
//CloseAll is a self generated function to close all windows
CloseAll();
//Select the Images folder, the macro will use the parent folder (dirgeneral) of the images folder as the place to save other folders and files generated
dirimage= getDirectory(“Please, select the images folder”);
dirgeneral=File.getParent(dirimage);
//generation of ROIs and Processed Images folders
roiFolder=dirgeneral+”/ROIs/”;
ProccFolder=dirgeneral+”/Processed/”;
if (File.exists(roiFolder)==false) File.makeDirectory(roiFolder);
if (File.exists(ProccFolder)==false) File.makeDirectory(ProccFolder);
//Checking of the existence of a Parameters file (ROIGParameters.txt) and if does not exists opens a Parameters menu to select ROIs names and visualization parameters
if (File.exists(dirgeneral+”/ROIGParameters.txt”)==false){
          n=getNumber(“How many ROI types do you want to define?”, 1);
          tiporoi=newArray(n);
          //Parameters menu
          Dialog.create(“Parameters Menu”);
          for(i=1; i<=n; i++) Dialog.addString(“Name for ROI type “+ i+ “:”,”ROI” +i);
          Dialog.addNumber(“Select the number of the channel you will use to draw contours”, 1);
          Dialog.addCheckbox(“Enhance contrast”, true);
          Dialog.show();
          //Variables collection from paramters menu
          for(i=1; i<=n; i++) tiporoi[i-1]=Dialog.getString();
          chan=Dialog.getNumber();
          enh=Dialog.getCheckbox();
          //Log register of ROIs Parameters
          print (“ROI types: \n”+n);
          print(“TIPO ROIs:”);
          for (i=0; i<tiporoi.length; i++) print(tiporoi[i]);
          print(“Channel:”);
          print(chan);
          print (“Enhance contrast:”);
          print(enh);
          selectWindow(“Log”);
          saveAs(“Text”, dirgeneral+”/ROIGParameters.txt”);
          selectWindow(“Log”);
          run(“Close”);
}
//Loading Parameters from ROIGParameters.txt file
else{
          param=File.openAsString(dirgeneral+”/”+”ROIGParameters.txt”);
          Vparam=split(param, “\n”);
          n=parseFloat(Vparam[1]);
          tiporoi=newArray(n);
          for (i=0; i<n; i++) tiporoi[i]=Vparam[3+i];
          chan=Vparam[n+4];
          enh=Vparam[n+6];
}
//Activation of drawing loop
previous=““;
*ImageLoad*(dirimage, roiFolder, ProccFolder);
//Function to find files inside the images folder
function *ImageLoad*(path, roipath, propath){
       folders=getFileList(path);
       for (i=0; i<folders.length; i++){
                if (File.isDirectory(path+folders[i])){
                          previous=substring(folders[i], 0, lengthOf(folders[i])-1);
                          //generation of group folder inside ROIs folder
                          roiFolder2=roipath+previous+”/”;
                          if (File.exists(roiFolder2)==false) File.makeDirectory(roiFolder2);
                          ProccFolder2=propath+previous+”/”;
                          if (File.exists(ProccFolder2)==false) File.makeDirectory(ProccFolder2);
                          *ImageLoad*(path+folders[i], roiFolder2, ProccFolder2);
             }
             else DrawRoi(path+folders[i], roiFolder2, ProccFolder2);
       }
}
//Function to Draw Rois
function DrawRoi(imagepath, roipath, propath){
       //Opening of image
       run(“Bio-Formats Importer”, “open=[“+imagepath+”] color_mode=Grayscale open_files view=Hyperstack stack_order=XYCZT”);
       name=File.nameWithoutExtension;
       if (enh==true) run(“Enhance Contrast”, “saturated=0.35”);
       run(“Brightness/Contrast…”);
       //call to the function to manually create ROIs
       rs=1;
       do{
               *DrawPol*(rs);
               sameimage=getBoolean(“Do you want to generate more ROIs in this image?”);
               rs++;
       }while (sameimage==true);
       //Closing and moving them to processed folder
       run(“Close All”);
       File.rename(imagepath, propath+folders[i]);
}
//function to draw ROIs in one particular image
function *DrawPol*(rs){
        //ROIs loop
        for (q=0; q<tiporoi.length; q++){
               roiFolder3=roipath+tiporoi[q]+”/”;
               //drawing tool
               setTool(“polygon”);
               do{
                         //Allowed user interaction
                         waitForUser(“Please, draw ROIs “+tiporoi[q]+”, and add them to the ROI
Manager by clicking t”);
                         //Rois checking
                         numberrois=roiManager(“count”);
                         if (numberrois==0){
                                 waitForUser(“YOU DID NOT DRAW ANY ROI”);
                                 cont=getBoolean(“Do you want to continue with next
ROI/Image?”);
                         }
                         else {
                                 roiManager(“Show All”);
                                 cont=getBoolean(“Do you want to save this/these ROIs?”);
                }
         }while (cont==false);
         numberrois=roiManager(“count”);
         if (numberrois>0 && cont==true){
                         roiManager(“Deselect”);
                         //Generation of folders for each type of ROI inside group and ROIs
folders
                         if (File.exists(roiFolder3)==false) File.makeDirectory(roiFolder3);
                         //Saving of ROIs
                         roiManager(“Save”, roiFolder3+name+rs+”.zip”);
                         roiManager(“reset”);
         }
     }
}
//This function closes all windows
function CloseAll(){
        run(“Close All”);
        list = getList(“window.titles”);
   for (i=0; i<list.length; i++){
        winame = list[i];
        selectWindow(winame);
        run(“Close”);
   }
}*
~~~

#### 3.4.3 Step by step guide of use for VolumestJ

1. After running the macro, a window with the Copyright License disclaimer will appear. To continue just click OK (more information about this License may be found by clicking the Help button).
2. A new window will prompt, asking the user whether she/he wants to analyze more than one section *per* image (Figure 6A). Click Yes if you generated ROIs for more than one section *per* image; otherwise, click No. The Cancel tab will end the macro.
3. A new window will be displayed where the user will be asked to select the directory that contains the Processed and ROIs folders. The organization of folders is depicted in Figure 5. We recommend maintaining the organization of folders generated after creating ROIs with the macro *VolumestJ_ROIsGenerator_v1.2.ijm*.
4. The macro will automatically open the first group of images (those from the first specimen, sample, or object) and their corresponding ROIs. The macro will analyze ROI areas, perimeters, and pixels (as minimum elements of the counting grid). Then, the macro will generate tables with the measurements and save them as .xls files in the Results folder (generated in the general directory previously selected).
5. The menu “Image Data” will be prompted (Figure 6B) where the user should provide three relevant parameters: a. the sections interval (e.g., if one of each 6, 1/6, sections were selected indicate 6), b. the thickness of the sections using the same units in which the images are calibrated, and c. the order of the sections. To indicate the order of sections, the user should select in each box the adequate number. Please, check the red numbers that appear in the image to identify the ROIs correctly. If one of the sections or images needs to be excluded from the analysis, select the X in the corresponding box, but in this case, please consider that no missing numbers are admitted except for the last ones (i.e., the sequence of selections 1, 2, X, 5, 4 will not be valid, the corresponding valid one is 1, 2, X, 4, 3). As previously indicated, organizing the sections in their correct order is relevant when estimating volumes using the TCS method. After choosing the adequate parameters, click OK and the macro will continue. The Cancel tab will end the macro.
6. The macro will estimate volumes using Cavalieri’s and TCS methods. A new table (Volumes) containing volume data and m0 and m1 coefficients of error (m0 and m1 CE) will be generated. Then, the macro will give some time to the user to check the table and save it (Figure 6C). At this point, it is recommended not to close the Volumes table, as it will be saved in .xls format inside the Results folder once all the images have been analyzed. By clicking OK, the macro will repeat steps 4 to 6.

**Figure 6.**
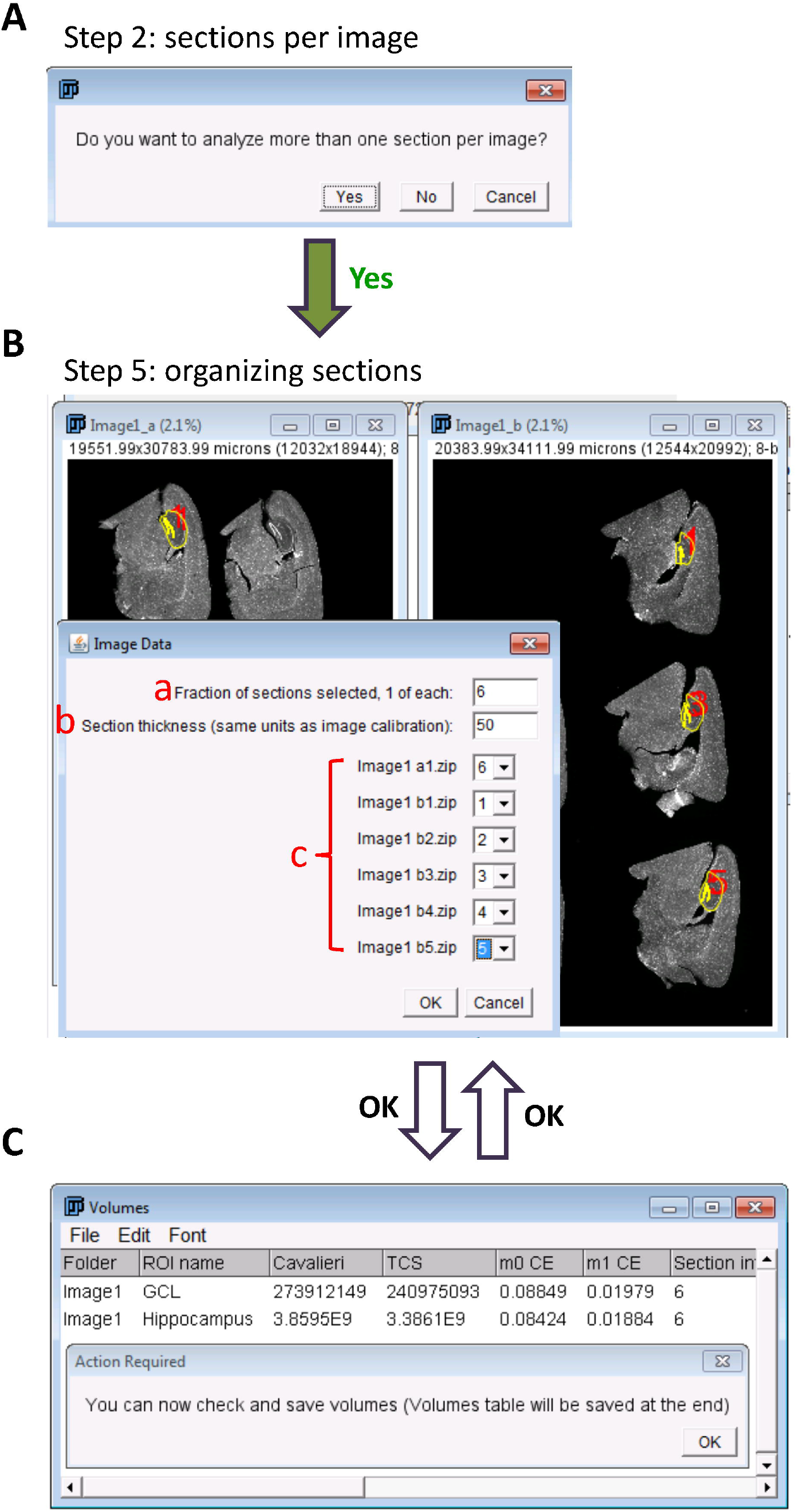
Relevant steps for volume estimation with VolumestJ. **A**. After accepting the copyright, VolumestJ will ask the user whether the images contain more than one section. By clicking Yes or No, the macro will analyze the first group of images and corresponding ROIs, create the Results folder (Figure 5), and save pixel, perimeter, and area tables in .xls format. Then, the macro will use the data obtained to estimate volumes and CEs, but it will need extra information provided by the user in step 5 (**B**). **B**. The macro will show all the images and ROIs used to analyze areas and perimeters, and a menu window will pop up. In the menu window, the user must indicate: a. the section interval as an entire number (eg. if the section interval is 1 of each 6, 1/6, the value should be 6), b. the thickness of the sections in the same units in which the image is calibrated, and c) the order of the sections (this is especially relevant for the estimation of volumes with the TCS method). If one of the sections will not be used, this should be indicated by selecting the X in the corresponding selection box of c. It should be considered that no missing numbers are allowed except for the last ones. **C**. After selecting parameters and clicking OK in step 5 (**B**), the macro will estimate volumes and CEs using the equations specified in this book chapter and then show the results in a table window named Volumes. The user will be allowed to check the results; please do not close the table, as it will not be saved until the macro ends (all the images and ROIs have been analyzed). By clicking OK in the Action Required window, the macro will analyze the next group of images pausing again at step 5.

~~~
**Macro code 5**
*// VolumestJ_v6.1.ijm
/*
   VolumestJ is an ImageJ macro developed to estimate volumes from pre-generated ROIs and
   microscopy images,
   Copyright (C) 2020 Jorge Valero Gómez-Lobo
   VolumestJ is free software: you can redistribute it and/or modify
   it under the terms of the GNU General Public License as published by
   the Free Software Foundation, either version 3 of the License, or
   (at your option) any later version.
   VolumestJ is distributed in the hope that it will be useful,
   but WITHOUT ANY WARRANTY; without even the implied warranty of
   MERCHANTABILITY or FITNESS FOR A PARTICULAR PURPOSE. See the
   GNU General Public License for more details.
   You should have received a copy of the GNU General Public License
   along with this program. If not, see <http://www.gnu.org/licenses/>.
*/
//This macro has been developed by Dr Jorge Valero (jorge.valero@achucarro.org).
//If you have any doubt about how to use it, please contact me.
//License
Dialog.create(“GNU GPL License”);
Dialog.addMessage(“VolumestJ Copyright (C) 2020 Jorge Valero Gomez-Lobo.”);
Dialog.setInsets(10, 20, 0);
Dialog.addMessage(“VolumestJ comes with ABSOLUTELY NO WARRANTY; click on help button for details.”);
Dialog.setInsets(0, 20, 0);
Dialog.addMessage(“This is free software, and you are welcome to redistribute it under certain conditions; click on help button for details.”);
Dialog.addHelp(“http://www.gnu.org/licenses/gpl.html”);
Dialog.show();
//CloseAll is a self generated function to close all windows;
CloseAll();
slicesuse=getBoolean(“Do you want to analyze more than one section per image?”);
run(“Set Measurements…”, “area perimeter redirect=None decimal=3”);
//Select the Images folder, the macro will use the parent folder (dirgeneral) of the images folder as the place to save other folders and files generated
generalDir= getDirectory(“Please, select general folder (it should contain the ROIS and Processed folders)”);
if (File.exists(generalDir+”Results/”)==false) File.makeDirectory(generalDir+”Results/”);
//Obtain image areas per group;
groups=getFileList(generalDir+”ROIs/”);
for (i=0; i<groups.length; i++){
                images=getFileList(generalDir+”Processed/”+groups[i]);
                ROIs=getFileList(generalDir+”ROIs/”+groups[i]);
                //Create tables for areas, perimeters and pixels
                tablearray=newArray(ROIs.length+1);
                tablearray[0]=“Image name”;
                for (ii=1; ii<tablearray.length; ii++) tablearray[ii]=substring(ROIs[ii-1], 0, lengthOf(ROIs[ii-1])-1);
                final=lengthOf(groups[i])-1;
                tabname=substring(groups[i], 0, final);
                TableCreator(“Areas”+tabname, tablearray);
                TableCreator(“Perimeters”+tabname, tablearray);
                TableCreator(“Pixels”+tabname, tablearray);
                //Open and measure image areas
                roisArr=newArray();
                for (ii=0; ii<images.length; ii++){
                               run(“Bio-Formats Importer”,
“open=[“+generalDir+”Processed/”+groups[i]+images[ii]+”] color_mode=Grayscale open_files view=Hyperstack stack_order=XYCZT”);
                               name=File.nameWithoutExtension();
                               rename(name);
                               run(“Enhance Contrast”, “saturated=0.35”);
                               sameimage=false;
                               sectioncounter=1;
                               do{
                                          //ROIs arrays to populate tables
                                          RAreas=newArray(ROIs.length+1);
                                          RPerim=newArray(ROIs.length+1);
                                          RPix=newArray(ROIs.length+1);
                                          RAreas[0]=name+sectioncounter;
                                          RPerim[0]=name+sectioncounter;
                                          RPix[0]=name+sectioncounter;
                                          sameimage=false;
                                          //Open and measure ROIs
                                          for(iii=0; iii<ROIs.length; iii++){
                                                      if
(File.exists(generalDir+”ROIs/”+groups[i]+ROIs[iii]+name+sectioncounter+”.zip”)){
                                                                If(roisArr.length>0){
                                                                        if
(name+sectioncounter+”.zip”!=roisArr[roisArr.length-1]){
                 roisArrtemp=newArray(name+sectioncounter+”.zip”);
                                                                        roisArr=Array.concat(roisArr,
roisArrtemp);
                                     }
                            }
                            else {
           roisArrtemp=newArray(name+sectioncounter+”.zip”);
                                     roisArr=Array.concat(roisArr, roisArrtemp);
                            }
                            roiManager(“open”,
generalDir+”ROIs/”+groups[i]+ROIs[iii]+name+sectioncounter+”.zip”);
                            n=roiManager(“count”);
                            if(n>1){
                                  roiManager(“Combine”);
                                  roiManager(“Add”);
                                  roiManager(“Select”, n);
                                  SliceNumber(sectioncounter);
                                  roiManager(“Select”, n);
                           }
                           else{
                                  roiManager(“Select”, 0);
                                  SliceNumber(sectioncounter);
                                  roiManager(“Select”, 0);
                           }
                           Overlay.addSelection
                           roiManager(“measure”);
                           RAreas[iii+1]=getResult(“Area”, 0);
                           RPerim[iii+1]=getResult(“Perim.”, 0);
                           getRawStatistics(nPixels, mean, min, max, std,
histogram);
                           RPix[iii+1]=nPixels;
                           roiManager(“reset”);
                           selectWindow(“Results”);
                           run(“Close”);
                           selectWindow(name);
                           run(“Select None”);
                           if (slicesuse==true){
                                      sectioncountertemp=sectioncounter+1;
                                      for(iv=0; iv<ROIs.length; iv++) {
      if(File.exists(generalDir+”ROIs/”+groups[i]+ROIs[iv]+name+sectioncountertemp+”.zip”))
{
                                                            sameimage=true;
                                                            iv=ROIs.length+1;
                                                 }
                                        }
                               }
                        }
              }
              //Populate tables
              TablePrinter(“Areas”+tabname, RAreas);
              TablePrinter(“Perimeters”+tabname, RPerim);
              TablePrinter(“Pixels”+tabname, RPix);
              sectioncounter++;
       }while (sameimage==true);
   }
   //Save tables
   SaveTable(“Areas”+tabname, generalDir+”Results/”);
   selectWindow(“Areas”+tabname);
   run(“Close”);
   SaveTable(“Perimeters”+tabname, generalDir+”Results/”);
   selectWindow(“Perimeters”+tabname);
   run(“Close”);
   SaveTable(“Pixels”+tabname, generalDir+”Results/”);
   selectWindow(“Pixels”+tabname);
   run(“Close”);
   do{
                      Table.open(generalDir+”Results/”+”Areas”+tabname+”.xls”);
   }while(isOpen(“Areas”+tabname+”.xls”)==false);
   do{
                      Table.open(generalDir+”Results/”+”Perimeters”+tabname+”.xls”);
   }while(isOpen(“Perimeters”+tabname+”.xls”)==false);
   do{
                      Table.open(generalDir+”Results/”+”Pixels”+tabname+”.xls”);
   }while(isOpen(“Pixels”+tabname+”.xls”)==false);
                      //Show images
                      run(“Tile”);
                      do{
                                repeat=false;
                                //Parameters and order of the images Menu
                                Dialog.create(“Image Data”);
                                Dialog.addNumber(“Fraction of sections selected, 1 of each: “,6);
                                Dialog.addNumber(“Section thickness (same units as image calibration):
“, 50);
                                if (slicesuse==true)images=Array.copy(roisArr);
                                it=newArray(images.length+1);
                                it[0]=“X”;
                                for (ii=1; ii<it.length; ii++) it[ii]=““+ii+”“;
                                for (ii=0; ii<images.length; ii++) Dialog.addChoice(images[ii], it, it[ii+1]);
                                Dialog.show();
                                //Menu values
                                run(“Close All”);
                                fraction=Dialog.getNumber();
                                thickness=Dialog.getNumber();
                                order=newArray(images.length);
                                for (ii=0; ii<images.length; ii++){
                                              choice=Dialog.getChoice();
                                              if (choice!=“X”) order[ii]=parseFloat(choice);
                                              else order[ii]=choice;
                                }
                                orderWO=Array.deleteValue(order, “X”);
                                orderWO=Array.sort(orderWO);
                                for (ii=0; ii<orderWO.length; ii++){
                                              if (orderWO[ii]!=ii+1){
                                                         repeat=getBoolean(“An error in the sequence of images has occured or a section is missing and volumes cannot be calculated”, “Back to Menu”, “DO NOT analyze these images”);
                                                         if (repeat==false) ROIs=newArray();
                                                         ii=orderWO.length+1;
                                              }
                                }
                    } while(repeat==true);
                    //Organize array for volume calculation
                    for (ii=0; ii<ROIs.length; ii++){
                              columnName=substring(ROIs[ii], 0, lengthOf(ROIs[ii])-1);
                              selectWindow(“Areas”+tabname+”.xls”);
                              ArrArea=Table.getColumn(columnName);
                              selectWindow(“Perimeters”+tabname+”.xls”);
                              ArrPerim=Table.getColumn(columnName);
                              selectWindow(“Pixels”+tabname+”.xls”);
                              ArrPix=Table.getColumn(columnName);
                              selectWindow(“Areas”+tabname+”.xls”);
                              ArrNames=Table.getColumn(“Image name”);
                              //Counting non used sections
                              Xn=0;
                              for (iii=0; iii<order.length; iii++) if(toString(order[iii])==“X”) Xn++;
                              //Ordering sections values
                              sizeArr=order.length-Xn;
                              OrgAreas=newArray(sizeArr);
                              OrgPerim=newArray(sizeArr);
                              OrgPix=newArray(sizeArr);
                              OrgNames=newArray(sizeArr);
                              for (iii=0; iii<order.length; iii++) if(toString(order[iii])!=“X”){
                                      OrgAreas[order[iii]-1]=ArrArea[iii];
                                      OrgPerim[order[iii]-1]=ArrPerim[iii];
                                      OrgPix[order[iii]-1]=ArrPix[iii];
                                      OrgNames[order[iii]-1]=ArrNames[iii];
                              }
                              //Call Cavalieri function
                              CavVol=Cavalieri(OrgAreas, fraction, thickness);
                              //Call ConicalVol function
                              ConicVol=ConicalVol(OrgAreas, fraction, thickness);
                              //Call Gundersen Coefficient of Error function
                              ArrGCE=GCE(OrgAreas, OrgPerim, OrgPix, fraction, thickness);
                              //Name of images used in order
                              ImOrd=““;
                              for(rr=0; rr<OrgNames.length; rr++) ImOrd= ImOrd+toString(rr+1)+”)
                              “+OrgNames[rr]+” “;
                              if (isOpen(“Volumes”)==false) VolTab();
                              tablearray=newArray(tabname, columnName, CavVol, ConicVol, ArrGCE[0], ArrGCE[1], fraction, thickness, ImOrd);
                              TablePrinter(“Volumes”, tablearray);
           }
           CloseAllEx(“Volumes”);
           waitForUser(“You can now check and save volumes (Volumes table will be saved at the end)”);
}
//Save Volume Table
if (isOpen(“Volumes”)) SaveTable(“Volumes”, generalDir+”Results/”);
//Cavalieri estimations
function Cavalieri(Areas, sectinterval, thickness){
            sumAr=0;
            n=Areas.length;
            //Summations
            for(i=0; i<n; i++) sumAr=sumAr+Areas[i];
            //Volume estimation
            V=sectinterval*thickness*sumAr;
            return(V);
}
//Conical volumes estimations
function ConicalVol(Areas, sectinterval, thickness){
            //Estimation of radius
            radius=newArray(Areas.length);
            for (i=0; i<Areas.length; i++) radius[i]=sqrt(Areas[i]/PI);
            //Estimation of distance between sections
            h=sectinterval*thickness;
            //Initial “virtual” section distance to first real section estimation
            h0=h/2;
            //Final “virtual” section distance to Final real section estimation
            hF=h/2;
            //Initial and final volumes estimations
            ini1=0;
            for (i=0; i<radius.length/2; i++){
                    if(radius[i]==0) ini1=i+1;
            }
            fin1=radius.length-1;
            for (i=radius.length-1; i>radius.length/2; i--){
                    if(radius[i]==0) fin1=i-1;
            }
            VolIni=PI*h0*pow(radius[ini1],2)/3;
            VolFinal=PI*hF*pow(radius[fin1],2)/3;
            SumVol=VolIni+VolFinal;
            //Total volume estimation
            for(i=0; i<radius.length-1; i++){
                    if (radius[i]>0 && radius[i+1]>0) {
            Voltemp=(PI*h*(pow(radius[i],2)+pow(radius[i+1],2)+(radius[i]*radius[i+1])))/3;
                         SumVol=SumVol+Voltemp;
                    }
           }
           return(SumVol);
}
//Gundersen Coefficient of Error estimation
function GCE(Areas, Perims, Pixels, sectinterval, thickness){
           ArrGCE=newArray(2);
           sumPix=0;
           n=Areas.length;
           sumshapeF=0;
           A=0;
           //mod is a variable that allows considering images without ROIs
           mod=0;
           //Summations
           for(i=0; i<n; i++){
                   sumPix=sumPix+Pixels[i];
                   A=A+pow(Pixels[i], 2);
                   if (Areas[i]>0) {
                         shapeF=Perims[i]/sqrt(Areas[i]);
                         sumshapeF=sumshapeF+shapeF;
                  }
                  else mod++;
          }
          n=n-mod;
          //Mean shape estimation
          meanShapeF=sumshapeF/n;
          //Variance due to noise estimation
          S2=0.0724*meanShapeF*sqrt(n*sumPix);
          //B and C estimations
          B=0;
          for(i=0; i<n-1; i++){
                     if (Pixels[i]>0 && Pixels[i+1]>0) B=B+(Pixels[i]*Pixels[i+1]);
          }
          C=0;
          for(i=0; i<n-2; i++){
                     if (Pixels[i]>0 && Pixels[i+2]>0) C=C+(Pixels[i]*Pixels[i+2]);
          }
          //Variance due to systematic random sampling
          Num=(3*(A-S2))-(4*B)+C;
          Varm0=Num/12;
          Varm1=Num/240;
          //m0 and m1 estimations
          TotalVarm0=S2+Varm0;
          TotalVarm1=S2+Varm1;
          //Gundersen coefficient of error estimations
          ArrGCE[0]=(sqrt(TotalVarm0))/sumPix;
          ArrGCE[1]=(sqrt(TotalVarm1))/sumPix;
          return ArrGCE;
          }
          //This function creates Volumes table
          function VolTab(){
                      tablearray=newArray(“Folder”, “ROI name”, “Cavalieri”, “TCS”, “m0 CE”, “m1 CE”, “Section interval”, “Section Thickness”, “Images order”);
                      TableCreator(“Volumes”, tablearray);
}
//This function closes all windows
function CloseAll(){
                      run(“Close All”);
                      list = getList(“window.titles”);
      for (i=0; i<list.length; i++){
                      winame = list[i];
                      selectWindow(winame);
                      run(“Close”);
      }
}
//This function closes all windows except one
function CloseAllEx(exception){
                      run(“Close All”);
                      list = getList(“window.titles”);
      for (i=0; i<list.length; i++){
                      winame = list[i];
                      if(winame!=exception){
                           selectWindow(winame);
                           run(“Close”);
                      }
         }
}
//This function generates tables
function TableCreator(tabname, tablearray){
                     run(“New… “, “name=[“+tabname+”] type=Table”);
                    headings=tablearray[0];
                    for (i=1; i<tablearray.length; i++)headings=headings+”\t”+tablearray[i];
                    print (“[“+tabname+”]”, “\\Headings:”+ headings);
}
//This function prints values in tables
function TablePrinter(tabname, tablearray){
                    line=tablearray[0];
                    for (i=1; i<tablearray.length; i++) line=line+”\t”+tablearray[i];
                    print (“[“+tabname+”]”, line);
}
//This function save tables
function SaveTable(tablename, dirRes){
                              selectWindow(tablename);
                              saveAs(“Text”, dirRes+tablename+”.xls”);
                    }
//This function place the text string1 in a pre-selected ROI
function SliceNumber(string1){
                   getPixelSize(unit, pixelWidth, pixelHeight); getDimensions(width, height, channels, slices, frames); x=getValue(“X”)/pixelWidth; y=getValue(“Y”)/pixelWidth;
                   run(“Select None”);
                   setFont(“SanSerif”, width/10, “antialiased”);
                             setColor(“red”);0
                   Overlay.drawString(““+string1+”“, x, y);
                   Overlay.show();
}*
~~~

### 3.5 Use of VolumestJ to detect sexual dimorphism of mouse hippocampus volume

We have tested the applicability of VolumestJ by checking the previously described dimorphism of the mouse hippocampus size [38]. The hippocampus has been divided into two regions attending to apparently different anatomical criteria: anterior vs posterior, dorsal vs ventral, and septal vs temporal [38–40]. However, despite the different nomenclature used, the anterior, dorsal and septal hippocampi almost comprise the same hippocampal regions and, thus, we will use here the nomenclature septal to refer to the anterior or dorsal portion of the hippocampus. This distinction is relevant as connectivity and functional differences have been described between the hippocampus’s septal and temporal regions [40, 41]. Here, we analyzed volumetric differences between the septal part of the GCL of the hippocampal dentate gyrus.

Our results indicate that the septal portion of the mouse GCL is significantly (*p*<0.05) larger in female than in male mice (Figure 7). This result agrees with a previous study indicating that the anterior portion of the mouse hippocampus is larger in females than in male mice [38]. VolumestJ generates a volume data file to compare Cavalieri’s and the TCS methods (Table 3). In this case, both methods allowed the detection of sexual dimorphism with a subtle difference in the scale of the change. Cavalieri’s method indicates that the female septal GCL is 1.14 times larger than the male one, while the TCS method shows a 1.15 times increase. Additionally, the variability of the TCS method is slightly smaller (female rse: 2.24%, male rse: 4.07%) than that obtained with Cavalieri’s method (female rse: 2.63%, male rse: 4.10%).

**Table 3.**
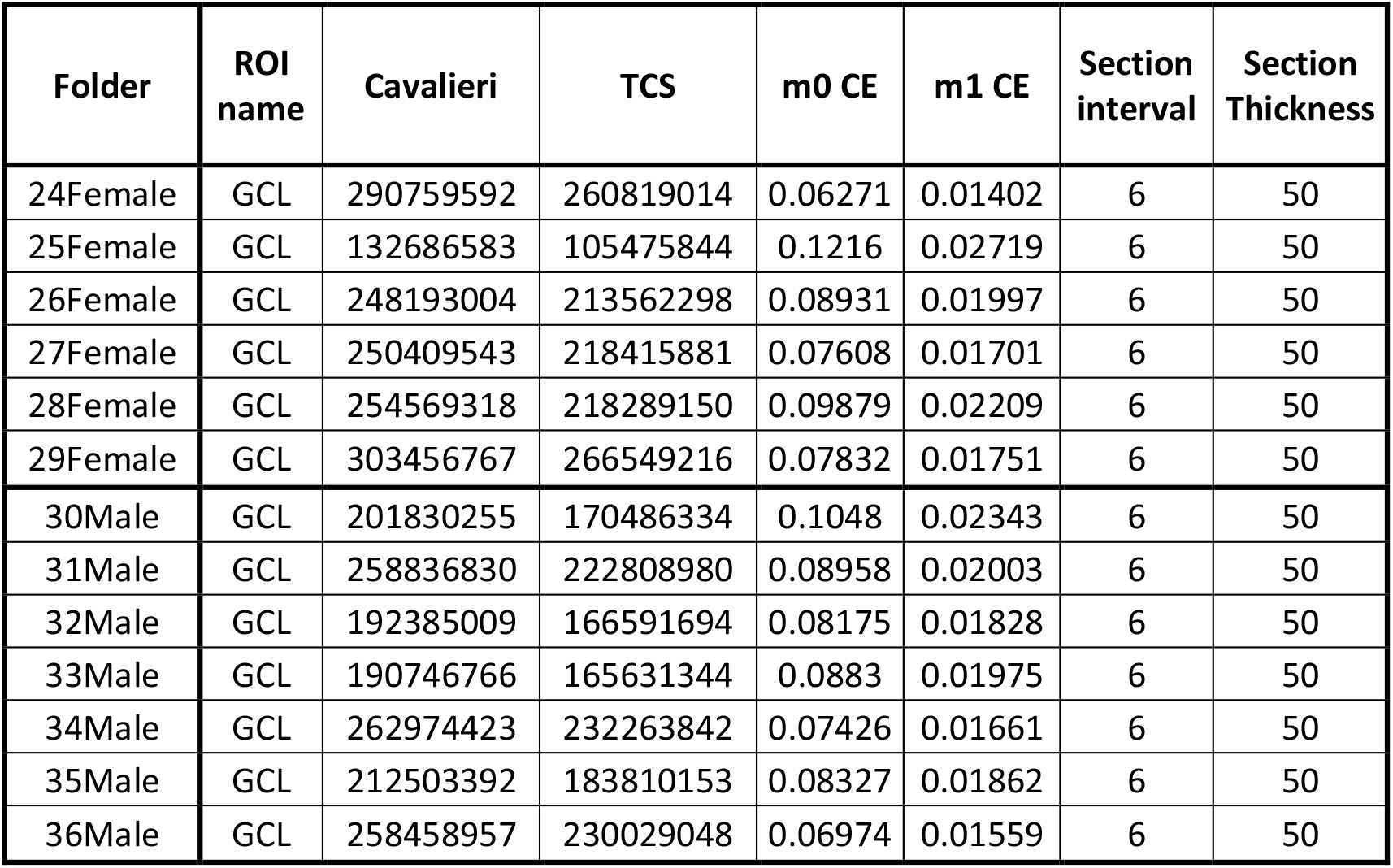
Data obtained using VolumestJ. The “Images order” column has been eliminated for reasons of space.

**Figure 7.**
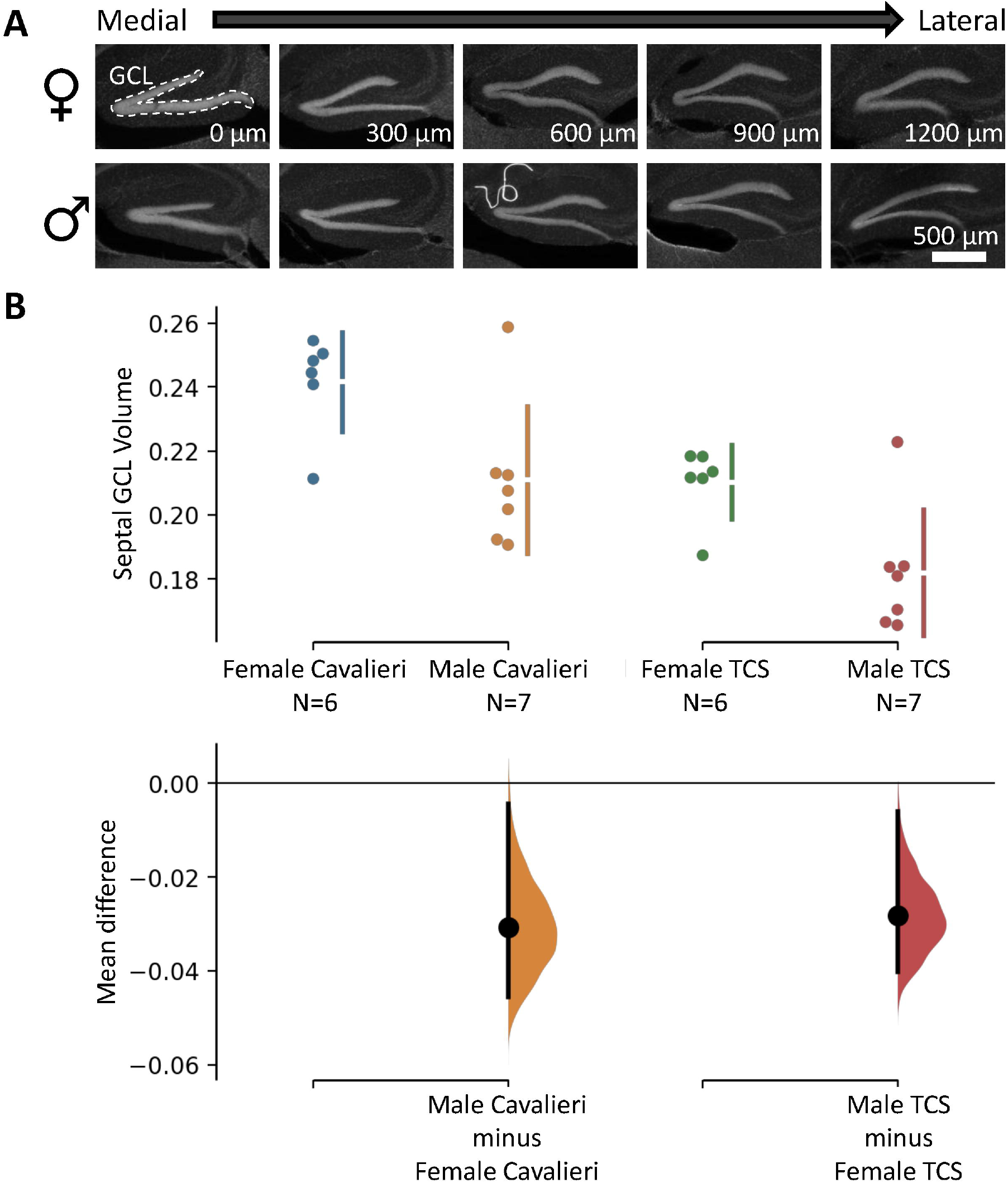
Analysis of sexual dimorphism in GCL volume using VolumestJ. **A**. Representative microscopy images of sagittal sections of female and male mouse brains showing the septal granule cell layer (GCL) of the hippocampal dentate gyrus. The numbers on the right-down corner indicate the relative distance to the first section (the most medial one). The scale bar in the bottom right image applies to all images. **B**. Cumming estimation plots showing the mean difference for 2 comparisons. Raw data are plotted on the upper axes; each mean difference is plotted on the lower axes as a bootstrap sampling distribution. Mean differences are depicted as dots, and the ends of the vertical error bars indicate 95% confidence intervals. The *p*-value of the two-sided permutation t-test is 0.0248 for Cavalieri’s method and 0.018 for the TCS method, thus indicating significantly larger volumes of the female septal GCL when compared with male mice. For each permutation *p*-value, 5000 reshuffles of the female and male labels were performed.

## 4 Conclusions

In this chapter, we have briefly revised the most commonly used method for volume estimation in science and proposed a new method (the TCS method) that has shown to be generally more accurate in artificially created shapes. We propose using pixel-associated measurements of areas as the best approach to estimate areas in the era of digital images. We have analyzed the advantages of our TCS method by challenging it to estimate different artificial objects of known volume and compare the results with those obtained with Cavalieri’s method. While the TCS method showed to be more robust in producing more accurate data when m1 CEs were lower than 0.03, in some specific conditions (check Figure 1) Cavalieri’s method may result in better estimations (geometrical prism shapes). Finally, we release a new ImageJ tool for Volume estimations (VolumestJ) which provides both Cavalieri’s and TCS method estimations. We hope this freely available software will facilitate volume estimations and extend the use of stereology.

## Acknowledgments

This work was supported by grants from the Spanish Ministry of Science and Innovation (https://www.ciencia.gob.es/) with FEDER funds (BFU2015-66689), a Tatiana Foundation project grant (P-048-FTPGB 2018), and the European Regional Development Fund (ERDF), through the COMPETE 2020 - Operational Programme for Competitiveness and Internationalization, and Portuguese national funds via FCT – Fundação para a Ciência e a Tecnologia (UIDB/04539/2020 and UIDB/04559/2020). Ferreiro E acknowledges FCT/ MCTES for her contract under the Scientific Employment Stimulus 2017 (CEECIND/00322/2017), Valero J salary was supported by Ikerbasque Basque Foundation for Science, Achucarro Basque Center for Neuroscience, Junta de Castilla y León/FEDER funds (SA0129P20) and the University of Salamanca; and Rodríguez-Iglesias N acknowledges the University of the Basque Country (UPV/EHU) for her predoctoral fellowship.

